# Hydrogen Sulfide modulates Flagellin-Induced Stomatal Immunity

**DOI:** 10.1101/2025.02.14.638267

**Authors:** Denise Scuffi, Rosario Pantaleno, Paula Schiel, Jan-Ole Niemeier, Alex Costa, Markus Schwarzländer, Ana M. Laxalt, Carlos García-Mata

## Abstract

Stomata are natural pores through which plants exchange gases with the environment, mainly carbon dioxide and oxygen required for photosynthesis and respiration, as well as water vapor through evapotranspiration. However, they also serve as entry points for microbial pathogens such as *Pseudomonas syringae* pv*. tomato* (*Pst*) bacteria. To prevent microbe invasion, guard cells detect pathogens-associated molecular patterns (PAMPs), including the bacterial peptide flagellin (flg22), triggering stomatal closure. This study identifies hydrogen sulfide (H_2_S) and its cytosolic source L-CYSTEINE DESULFHIDRASE 1 (DES1), as key players in stomatal immunity. We demonstrate that H_2_S and DES1 are involved in flg22- and bacterial-induced responses, including stomatal closure and modulation of reactive oxygen species (ROS) production. We have found that knock out mutants in *DES1* gene exhibits reduced susceptibility to *Pst* spray-inoculation and lower apoplastic and cytosolic H_2_O_2_ levels in response to flg22. Additionally, H_2_S independently induces cytosolic H_2_O_2_ levels in guard cells without requiring RBOHD activity. All together, these findings establish H_2_S and its source, DES1, as critical components of the stomatal immune response.

**One Sentence Summary:** H_2_S and DES1 actively participate in flg22-induced stomatal closure modulating apoplastic and cytosolic ROS production.

## INTRODUCTION

Plants are exposed to a wide variety of environmental challenges, including a diversity of pathogens. Stomata act as a hub of the exchange between the plant and its environment in the aerial part of land plants, because they enable and regulate the large fluxes of inorganic matter, i.e. water, carbon dioxide and oxygen that ultimately build and maintain the plant homeostasis. The size of the stomatal pore is regulated through variations of guard cell volume. Multiple stimuli, both environmental and endogenous, are sensed by plants and translated through a complex signalling network into redistribution of osmotically active solutes, resulting in the influx/efflux of water with the consequent change in cell volume. Since stomatal opening has direct impact on carbon fixation and water homeostasis, stomatal pore regulation is a central physiological process for the plant, and scales up to our global climate (Blatt, 2000; Schroeder et al., 2001; Kim et al., 2010; Qi et al., 2018).

A potential downside of “natural openings” in the epidermal layer of plants, such as stomata, hydathodes or wounds, is that they provide an entry point for many pathogens, including bacteria and some fungi. Upon contact with pathogens-associated molecular patterns (PAMPs), stomatal closure is triggered as an early defence response to avoid the infection of the endogenous tissues (Underwood et al., 2007; Agurla et al., 2014; Melotto et al., 2017). In Arabidopsis, flagellin and flg22 (a conserved peptide from flagellin that is recognized as a PAMP) are recognized by the membrane immune receptor, FLAGELLING SENSING 2 (FLS2), to induce PAMP-triggered immunity (PTI). This signalling event, among others, involves the production of superoxide (O_2_.^-^) via the activation of RESPIRATORY BURST OXIDATIVE HOMOLOGUE D (RBOHD) which then is converted by the superoxide dismutase to hydrogen peroxide (H_2_O_2_) in the apoplast; calcium (Ca^2+^) influx from the apoplast; the activation of mitogen activated-protein kinases (MAPKs) and stomatal closure (Bethke et al., 2012; Daudi et al., 2012; Kadota et al., 2014a; Li et al., 2014; Toum et al., 2016; Arnaud et al., 2017; Tian et al., 2019; Thor et al., 2020; Bjornson et al., 2021). The signalling triggered by PAMPs that leads to stomatal closure is referred to as stomatal immunity, one of the plant’s initial responses to pathogen attack to limit their entry into the leaf (Melotto et al., 2024). Some pathogens, such as *Pseudomonas syringae* pv. *tomato* DC3000 (*Pst* DC3000), have developed mechanisms to suppress stomatal immunity by “rewiring” guard cell signalling network through the production of phytotoxins like coronatine (COR), to reopen the stomata after 3h of infection (Melotto et al., 2006). Despite major advances in the understanding of the stomatal immunity signalling network, new actors and mechanisms of action are still emerging.

Hydrogen sulfide (H_2_S) acts as a gasotransmitter in many biological systems, including plants, where it plays a key role in guard cell signalling (Pantaleno et al., 2021). There is evidence that H_2_S modulates proteins by persulfidation, i.e. a posttranslational modification (PTM) of cysteine (Cys) residues in target proteins (Wang et al., 2021; Pantaleno and Scuffi., 2024). Formation of persulfides (RSSH) on Cys thiol moieties is reversible, giving rise to a molecular switch in cell signalling that may be linked to the modulation of molecular structure and/or activity of the modified protein. This can be observed in proteomes of persulfidated proteins from Arabidopsis leaf extracts, where some of the hits are modified in a reversible manner (Aroca et al., 2015; Aroca et al., 2017). Persulfidation is currently regarded as a mechanism that protects proteins from irreversible oxidation in persistent oxidative environments (Filipovic et al., 2018; Aroca et al., 2021). Plants synthesize H_2_S enzymatically from sulfur-containing amino acids in different subcellular compartments (Gotor et al., 2019). Among them, the first characterized source, L-CYSTEINE DESULFHYDRASE 1 (DES1), which degrades L-cysteine to produce pyruvate, ammonia and H_2_S, is considered a major cytosolic source (Álvarez et al., 2010). DES1-dependent H_2_S production participates in several physiological processes, including stomatal closure in response to different stimuli (Scuffi and García-Mata, 2021). For example, in abscisic acid (ABA)-dependent stomatal closure, guard cell specific *DES1* expression increases cytosolic H_2_S (Scuffi et al., 2014; Du et al., 2019; Zhang et al., 2019) which, in turn, persulfidates and modulates the activity of key enzymes of the guard cell signalling network such as OPEN STOMATA 1 (OST1) and RBOHD, resulting in stomatal closure (Chen et al., 2020; Shen et al., 2020). In addition, H_2_S was reported to inhibit inward-rectifying K^+^ channels and increase the cytosolic levels of other second messengers such as nitric oxide (NO), phosphatidic acid (PA) and cytosolic H_2_O_2_ specifically in guard cells (Scuffi et al., 2014; Papanatsiou et al., 2015; Scuffi et al., 2018). However, the above-mentioned responses are not necessarily linked to ABA signalling.

The involvement of H_2_S, and its enzymatic sources, in abiotic stress responses, has attracted the research attention of plant scientists. In contrast, the putative role of H_2_S in plant-pathogen interactions may have lagged behind. Fungal infection can boost H_2_S emission in different crop species through a process known as Sulfur Induced Resistance (SiR). SiR is intimately related to sulfur metabolism and to the nutrient status of the plant (Bloem, 2004; Bloem et al., 2007; Bloem et al., 2012; Vojtovič et al., 2021). In addition, bacterial pathogens, such as *Pst* DC3000, were reported to modulate plant H_2_S dynamics, leading to resistance (Shi et al., 2015). Furthermore, we have recently established that the mitochondrial source of H_2_S, β-cyanoalanine synthase (CAS-C1) participates in flg22-induced stomatal closure (Pantaleno et al., 2024b). Although, the involvement of H_2_S in stomatal immunity has been poorly investigated.

In this work we set out to understand the involvement of cytosolic DES1/H_2_S production in stomatal immunity and its relationship with other components of the PTI response, such as the second messengers H_2_O_2_ and Ca^2+^. To specifically dissect stomatal responses, we combined the genetic impairment of cytosolic H_2_S production with cell compartment-specific biosensing of responses in guard cell physiology. Our findings integrate cytosolic H_2_S into the stomatal signalling network.

## RESULTS

### Absence of DES1 affects Pseudomonas-induced stomatal closure and immunity

To investigate the role of H_2_S signalling in stomatal immunity we used *des1* mutant lines of Arabidopsis to abolish the main cytosolic H_2_S source. We first exploited this model to address the question of whether there is any involvement of H_2_S in stomatal movement in response to *Pst* DC3000 versus *Pst* DC3000 *hrcC^-^*, being the latter a strain with a defective type III secretion system as required for effector secretion. *Pst* DC3000 *hrcC^-^*represents a useful tool to study PTI responses without the interference of bacterial effectors (Hauck et al., 2003). Epidermal peels from the abaxial side of Arabidopsis leaves from wild type (Col-0) and *des1* mutant plants were isolated and incubated in opening buffer and subsequently treated with a bacterial suspension of *Pst* DC3000 (Figure 1 A) or *Pst* DC3000 *hrcC^-^* (Figure 1 B) for 1 h or 3 h. Stomata exposed to either of the bacterial suspensions close after 1 h and reopen after 3 h of treatment in Col-0 epidermal peels, as compared to the mock control. Although *des1* stomata exhibited lower stomatal aperture values with the mock treatment, we have previously shown they have the capacity to close under H_2_S-donor and H_2_O_2_ treatment (Scuffi et al., 2014; Scuffi et al., 2018). However, in contrast to Col-0, no difference in aperture was observed in stomata from *des1* plants among treatments, suggesting that DES1 is required for the full scale stomatal PTI response to *Pst* (Figure 1).

**Figure 1:**
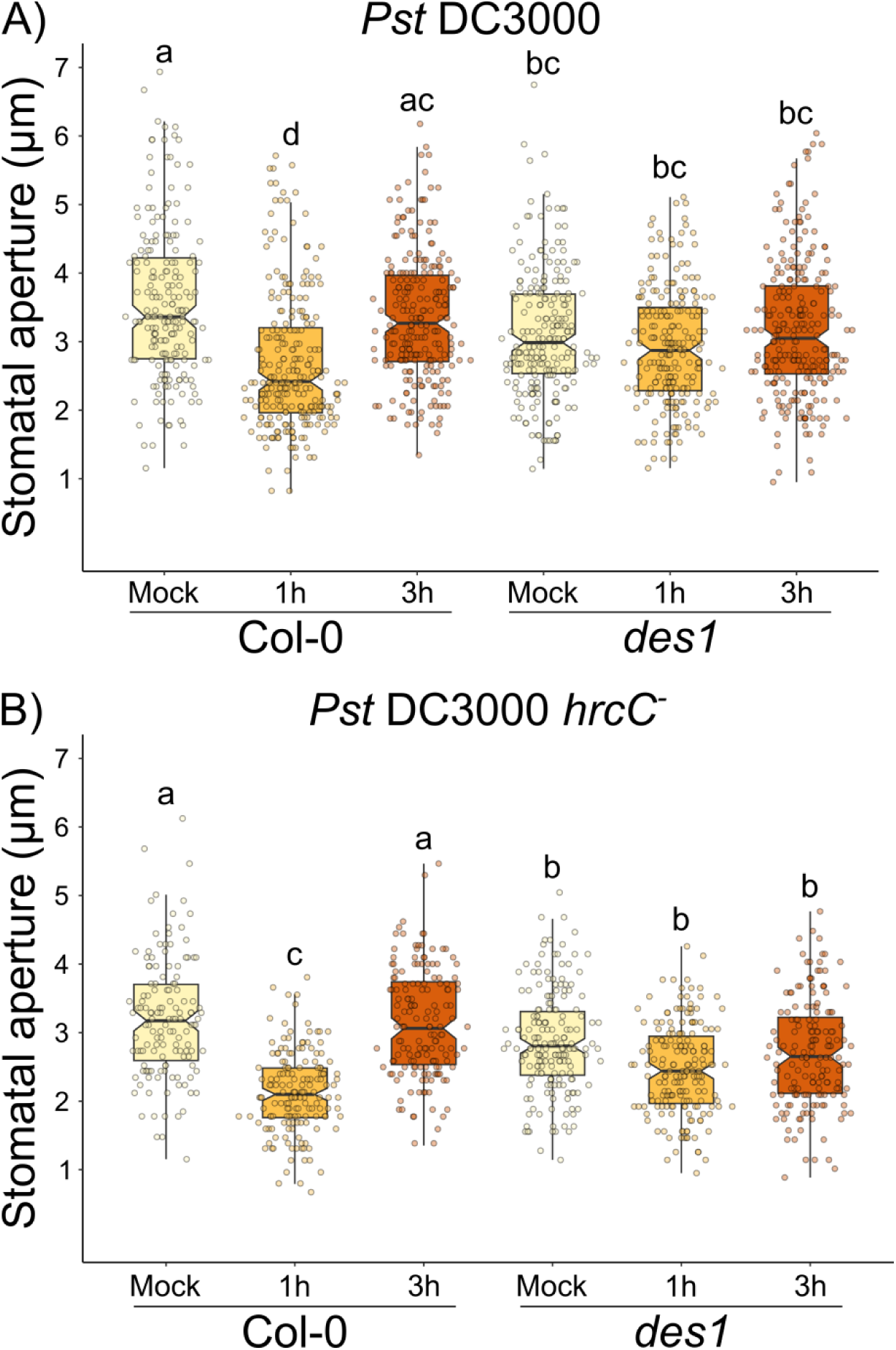
Absence of DES1 affects Pseudomonas-induced stomatal closure. Epidermal peels from 4- to 5-week-old wild type (Col-0) and *des1* (*des1*) mutant Arabidopsis plants were pre-incubated in opening buffer (5 mM MES pH 6.1, 50 mM KCl) for 3 h under light and subsequently treated with 24 h-grown bacteria (OD=0.1) *Pst* DC3000 (A) or *Pst* DC3000 *hrcC*^-^ mutant (B) in the same buffer under light for 1 h or 3 h. The values of stomatal aperture are expressed in microns (µm) and represented in box-plots where the box is bound by the 25^th^ and 75^th^ percentile, whiskers span 10^th^ and 90^th^ percentile, and the line in the middle is the median. The individual points represent each measurement. Data is from four independent experiments (Table S5). Different letters indicate statistical differences among treatments (Tukey’s Method, p-value < 0.05).

Given that there is no information on the involvement of DES1 and H_2_S in the response of leaf tissue to flg22, we asked how DES1/H_2_S may affect flg22-triggered stomatal signalling. To address this question, we isolated epidermal peels from Col-0 and *des1* leaves and treated them with 0, 10, 100 and 1000 nM of flg22. Flg22 induced stomatal closure in Col-0 epidermal peels in a concentration-dependent manner while this response was disrupted in *des1* stomata to the same flg22 concentrations (Figure 2 A). To determine if the impaired stomatal immunity of *des1* mutant is due to the reduction of H_2_S production, we challenged *des1* epidermal peels with flg22 together with the H_2_S donor, GYY4137. Figure 2 B shows that exogenous addition of H_2_S restores stomatal response to flg22 indicating that H_2_S is required for flg22-triggered stomatal immunity response (Figure 2 B). Finally, in order to see whether the response was specific to flg22, we tested other PAMP such as the bacterial elongation factor Tu (EF-Tu) peptide 18 (elf18), which consistently caused stomatal closure in Col-0 but not in *des1* mutant (Figure S1). Taken together, these results strongly suggest that DES1 participates in stomatal closure elicited by PAMPs.

**Figure 2:**
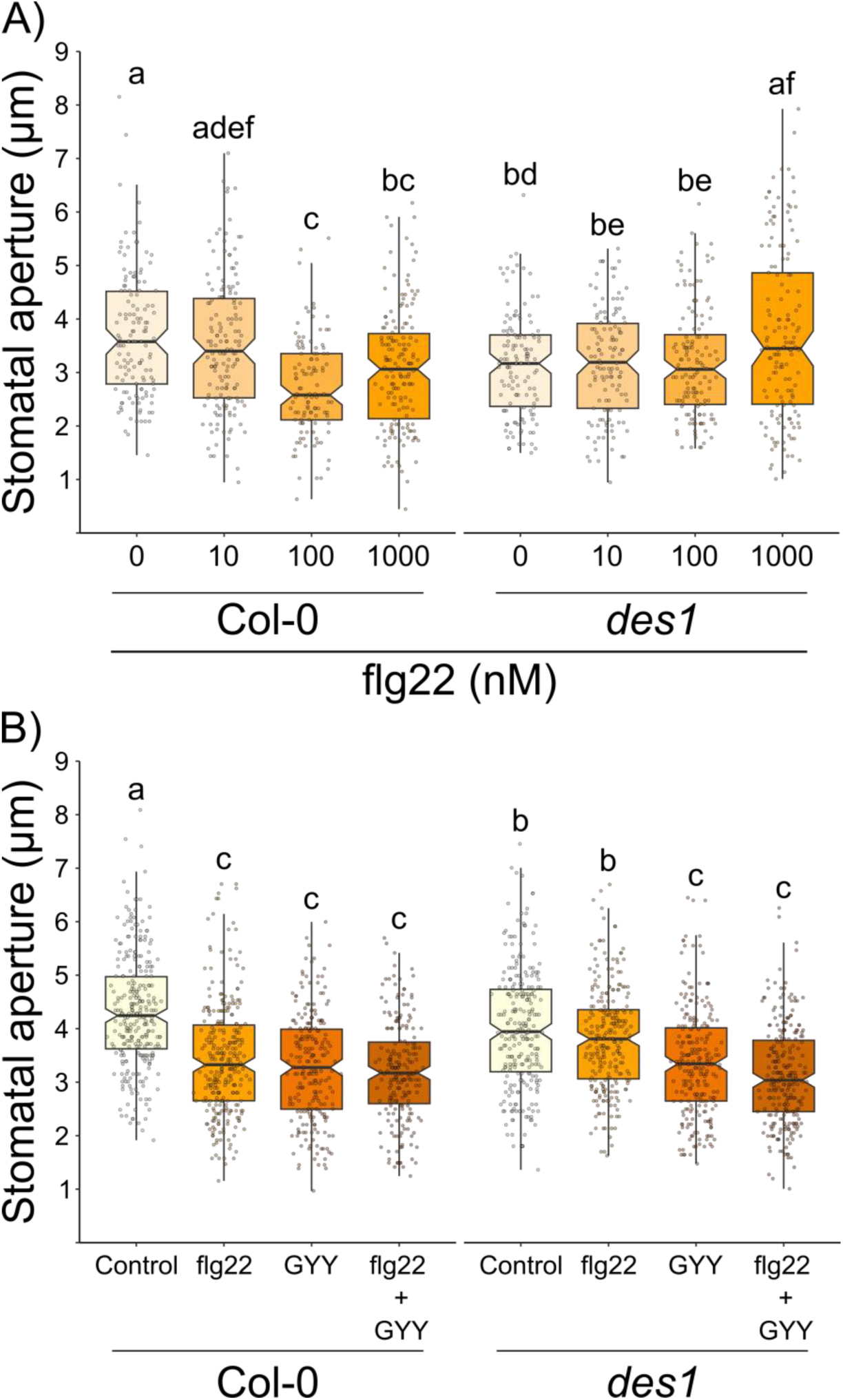
DES1 is required for flg22-induced stomatal closure response. Epidermal peels from 4- to 5-week-old wild type (Col-0) and *des1* (*des1*) mutant Arabidopsis plants were pre-incubated in opening buffer (5 mM MES pH 6.1, 50 mM KCl) for 3 h under light and subsequently treated for 90 min with 0, 10, 100 or 1000 nM flg22 (A), or with opening buffer (Control), 1 µM flg22, 100 µM of the H_2_S donor, GYY4137 (GYY) or flg22 + GYY4137 (B) in the same buffer under light. The values of stomatal aperture are expressed in microns (µm) and represented in box-plots where the box is bound by the 25^th^ and 75^th^ percentile, whiskers span 10^th^ and 90^th^ percentile, and the line in the middle is the median. The individual points represent each measurement. Data is from at least three independent experiments (Table S5). Different letters denote statistical differences among treatments (Tukey’s Method, p-value < 0.05).

Since DES1 participates in bacterial and elicitor-dependent stomatal closure, we inoculated *des1* mutant plants with *Pst* DC3000 *hrcC^-^* bacteria by spray, in order to study the susceptibility of these plants taking into account the participation of the stomata in this response. Although an increase in bacterial growth is observed at 72 hours after inoculation compared to 24 hours in both genotypes, *des1* mutant plants show less growth compared to the Col-0 plants indicating a lower susceptibility phenotype (Figure 3)

**Figure 3:**
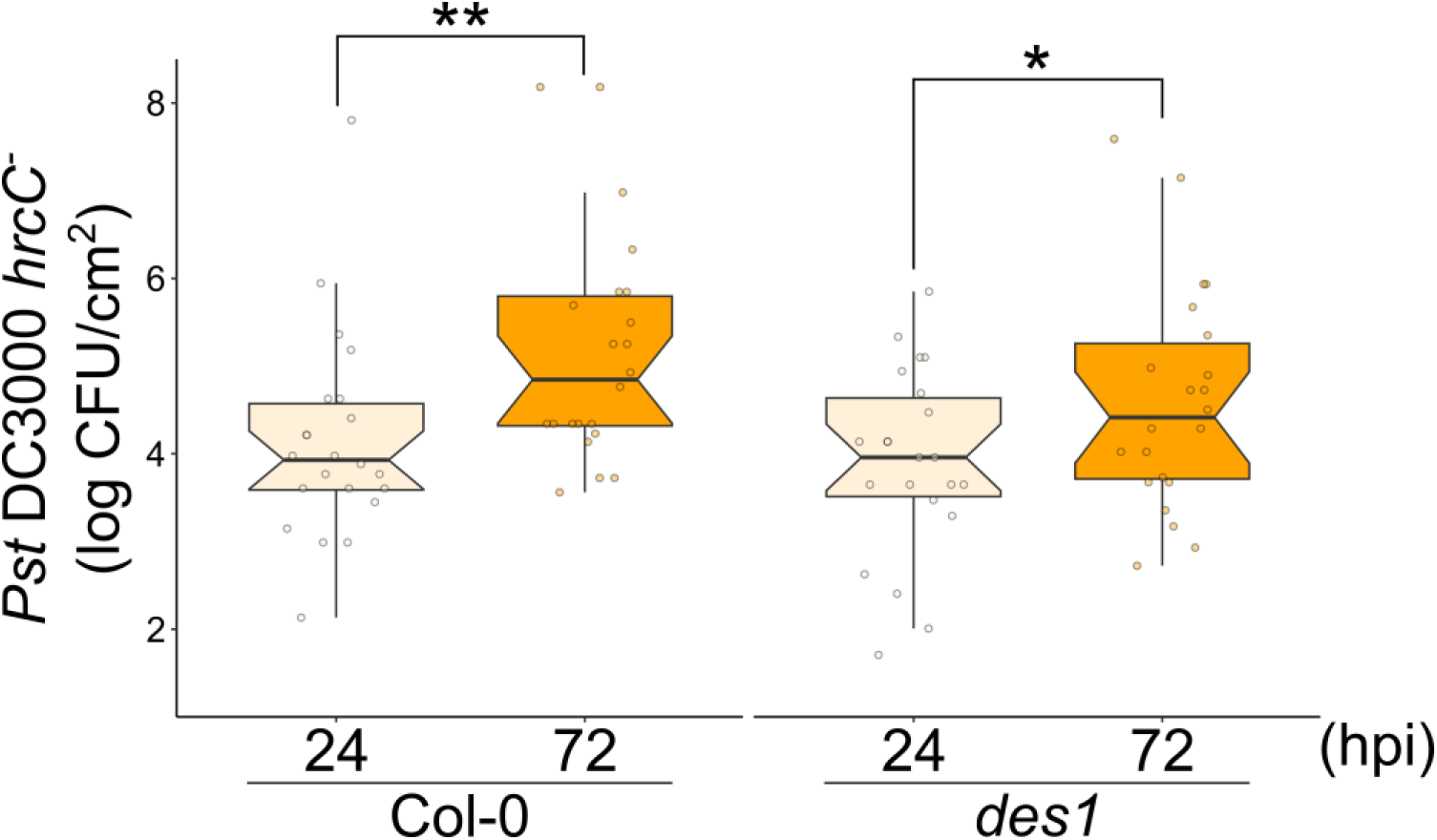
*des1* mutant plants are less susceptible to *Pst.* DC3000 *hrcC^-^* spray inoculation. 5-6 week-old Arabidopsis wild type (Col-0) and *des1* (*des1*) plants were surface-inoculated by spray with a *Pseudomonas syringae* pv. *tomato* DC3000 *hrcC^-^* (*Pst.* DC3000 *hrcC^-^*) suspension (OD = 0.2 (λ = 600 nm)). Bacterial growth was assessed in leaf discs at 24- or 72-hours post-inoculation (hpi), and the number of colony-forming units (CFU) per cm^2^ of leaf extracts was determined. Log CFU/cm^2^ were calculated and represented in box-plots where the box is bound by the 25^th^ to 75^th^ percentile, whiskers span 10^th^ to 90^th^ percentile, and the line in the middle is the median. The individual points represent each measurement (Table S5). Asterisks denote statistical differences between treatment (Paired samples *t*-test, P-value <0.05 = (*), P-value <0.01 = (**)).

### DES1 is involved in flg22-induced apoplastic H_2_O_2_ production

Flg22 perception triggers an apoplastic, RBOHD-dependent, ROS burst within the first minutes of the defense response (Felix et al., 1999; Kadota et al., 2014a). To assess the involvement of DES1 in the RBOHD-branch of the stomatal PTI signalling network, we assayed apoplastic ROS production by flg22-treated leaf discs as a readout using luminol-based detection method. Strikingly, the apoplastic ROS burst was attenuated in *des1* leaf discs as compared to Col-0 leaf discs (Figure 4), pinpointing a role for DES1 in the RBOHD-mediated redox signaling elicited by flg22.

**Figure 4:**
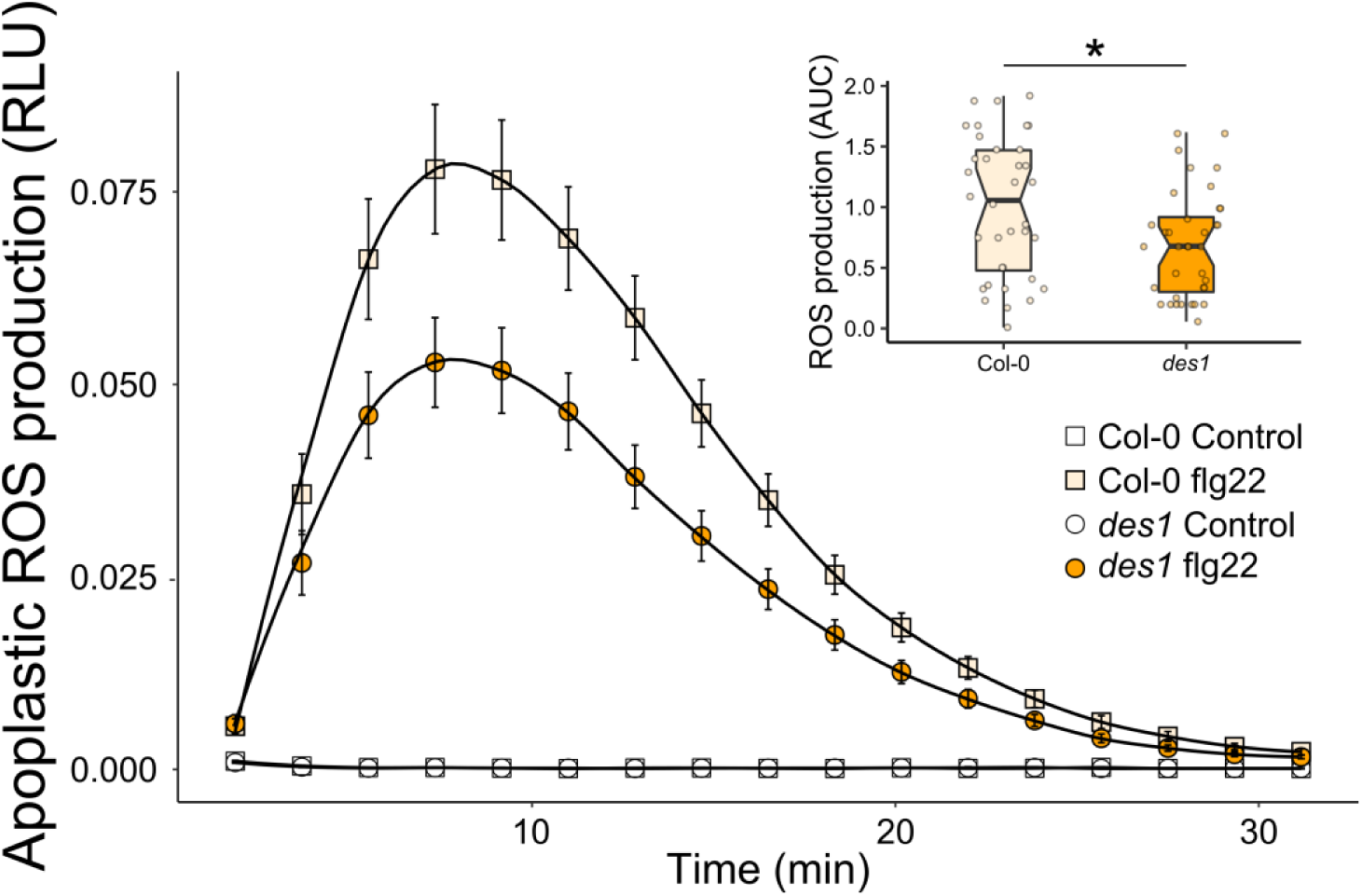
Flg22-induced apoplastic ROS burst is diminished in *des1* leaf discs. Leaf discs of 4- to 5-week-old wild type (Col-0) or *des1* (*des1*) mutant Arabidopsis plants were incubated with 100 nM flg22 and the ROS burst was measured with a luminol-based assay. The luminescence was recorded every 2 min for 30 min and expressed as relative light units (RLU). The curves show the mean± SE of 12 discs from three independent experiments (n=36) during time (Table S4). Inset: Total ROS production was calculated integrating areas under curve (AUC) from each leaf disc and represented in box-plots where the box is bound by the 25^th^ to 75^th^ percentile, whiskers span 10^th^ to 90^th^ percentile, and the line in the middle is the median. The individual points represent each measurement (Table S5). Asterisk denotes statistical differences between treatments (*t*-test, p-value 0.01)

### Flg22-induced cytosolic H_2_O_2_ signature is DES1 and H_2_S-dependent

After the rapid apoplastic ROS burst, flg22 induces an increase in cytosolic H_2_O_2_ which is required for stomatal closure induction (Toum et al., 2016; Rodrigues et al., 2017; Nietzel et al., 2019; Arnaud et al., 2023a). To study the role of DES1 in cytosolic H_2_O_2_ dynamics in response to flg22, we isolated epidermal peels from Col-0 and *des1* mutant Arabidopsis plants expressing the specific cytosolic H_2_O_2_ biosensor roGFP2-Orp1 (Nietzel et al., 2019) and floated them in opening buffer for at least 7 h to ensure full recovery from tissue injury-induced oxidation as we have previously described (Scuffi et al., 2018; Pantaleno et al., 2024a). Then, epidermal peels were treated with flg22 and the redox state of the cytosolic localised sensor in guard cells was determined. The first observation is that *des1* exhibits a higher oxidation state of the sensor under control conditions (Figure 5). Upon flg22 treatment, oxidation of roGFP2-Orp1 was induced in both Col-0 and *des1* background, indicating an increase in cytosolic H_2_O_2_ concentration. However, the magnitude of the response in *des1* was significantly lower compared with Col-0, (p-value < 2.2e-16, Wilcoxon test in Col-0 vs p-value = 3.668e-10, Wilcoxon test in *des1*) (Figure 5). Notably, flg22-dependent oxidation was abolished by pre-treatment with the H_2_S scavenger hypotaurine (HT) (Figure S2). Taken together, these results indicate that DES1 and, more broadly, endogenous H_2_S are required to induce cytosolic H_2_O_2_ production in guard cells in response to flg22.

**Figure 5:**
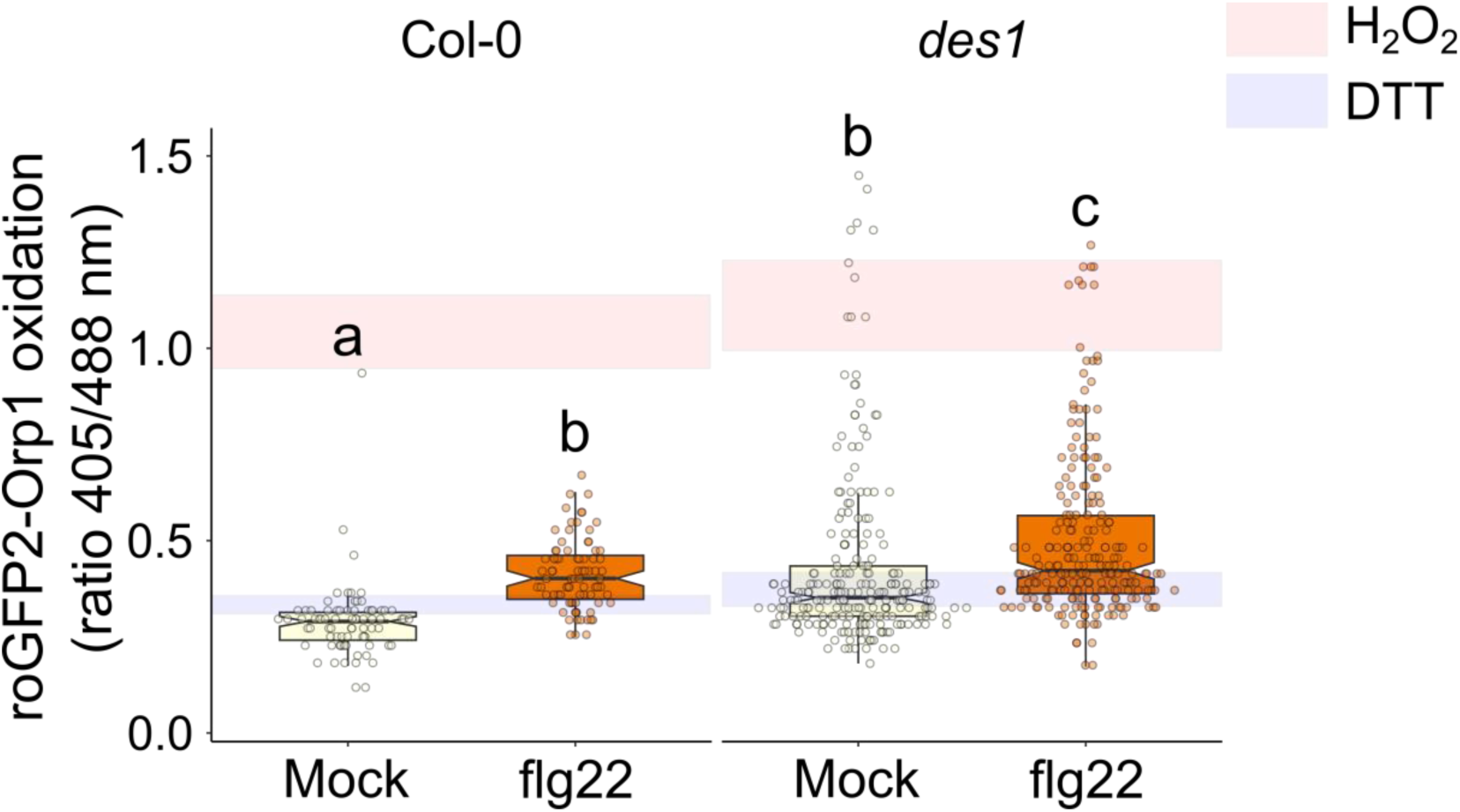
Flg22 requires DES1 to induce cytosolic H_2_O_2_ flux in guard cells. Epidermal peels from 4- to 5-week-old Col-0 or *des1* mutant plants expressing H_2_O_2_ specific biosensor roGFP2-Orp1 in the cytosol were incubated for 7-12 h in opening buffer (5 mM MES pH 6.1, 50 mM KCl). Then, were treated with opening buffer (Mock) or 1 µM flg22 (flg22) for 60-90 min. Moreover, epidermal peels were treated with 20 mM DTT or 10 mM H_2_O_2_ for 10 min to estimate the dynamic range of the sensor *in situ*. Values are expressed as the ratio of 405/488 nm and are represented in the box plots where the box is bound by the 25^th^ to 75^th^ percentile, whiskers span 10^th^ to 90^th^ percentile, and the line in the middle is the median. The individual points represent each measurement. Red and blue bands indicate the 25^th^ to 75^th^ percentile of maximum and minimum ratio values obtained for treatments with external H_2_O_2_ and DTT, to drive the sensor towards full oxidation and reduction. Data are from at least three independent experiments (Table S5). Different letters denote statistical differences among treatments (Tukey’s Method, p-value < 0.05)

### H_2_S-dependent cytosolic H_2_O_2_ rise in guard cells is not via RBOHD

We next asked the question of whether the H_2_S-dependent cytosolic H_2_O_2_ response is dependent on RBOHD or rather on a different branch of the PTI pathway. We generated lines expressing the cytosolic roGFP2-Orp1 sensor in the *rbohD* background (Torres et al., 2002) by crossing. Epidermal peels from Col-0 and *rbohD* expressing roGFP2-Orp1 were incubated in opening buffer and then treated with 100 µM of the H_2_S donor, GYY4137 for 15 minutes as previously established (Scuffi et al., 2018). GYY4137-derived H_2_S induced an increase in roGFP2-Orp1 oxidation in Col-0 guard cells. However, roGFP2-Orp1 oxidation of *rbohD* guard cells was even greater than that of wild type plants (Figure 6) demonstrating that RBOHD is not required for the cytosolic H_2_O_2_ response to H_2_S.

**Figure 6:**
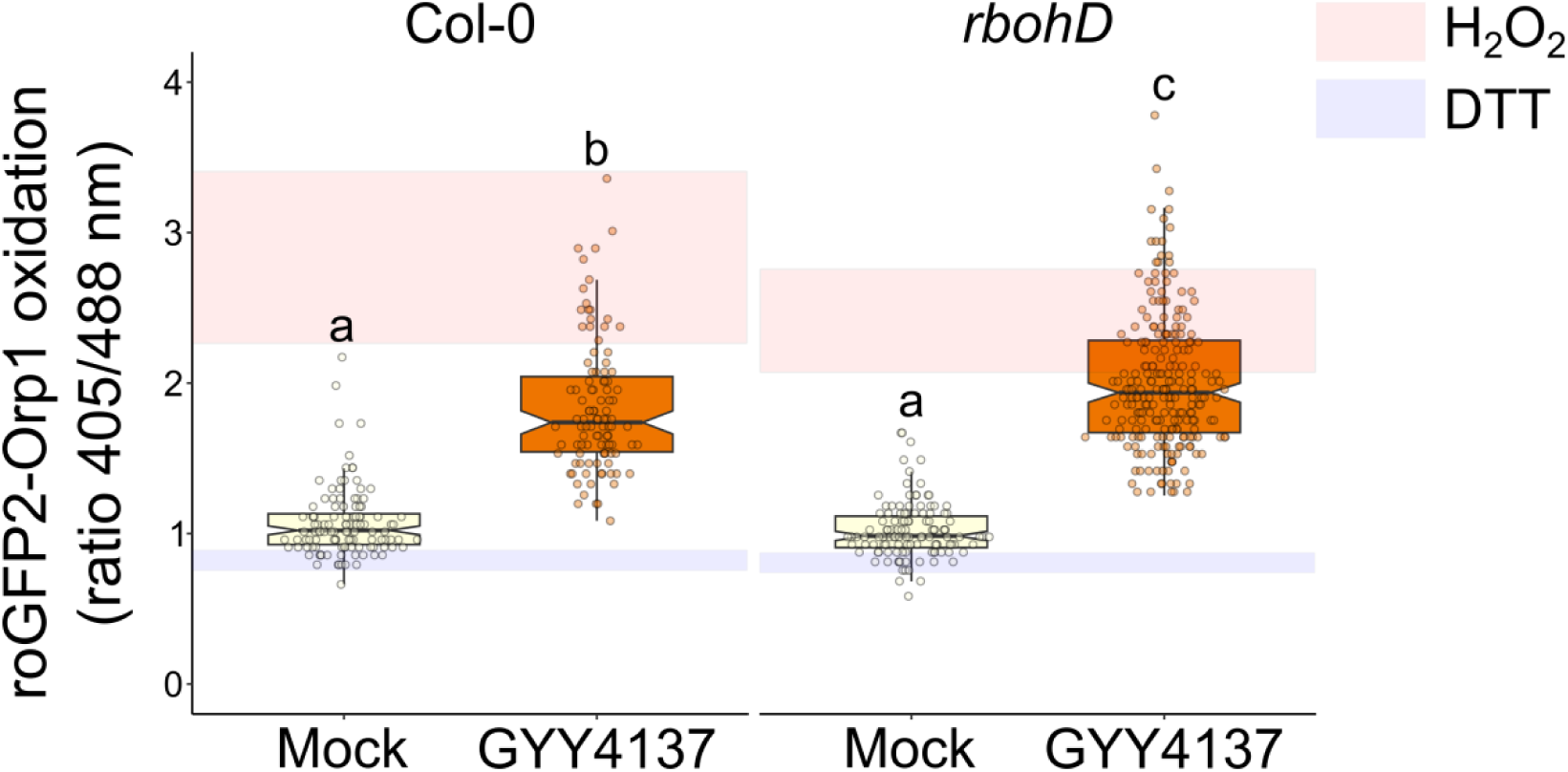
H_2_S-induced cytosolic roGFP2-Orp1 oxidation in guard cells does not require RBOHD. Epidermal peels from from 4- to 5-week-old Col-0 and *rbohD* Arabidopsis plants expressing the H_2_O_2_ biosensor, roGFP2-Orp1 in the cytosol were incubated in opening buffer (5mM MES pH 6.1, 50 mM KCl) for 7-12 h and then treated with Dimethyl sulfoxide (DMSO) 0.01% (v/v) (Mock) or with 100 µM of the H_2_S donor GYY4137 (GYY4137) for 15 min. Moreover, epidermal peels were treated with 20 mM DTT or 10 mM H_2_O_2_ for 10 min to estimate the dynamic range of the sensor *in situ*. Values are expressed as the ratio of 405/488 nm and are represented in the box plots where the box is bound by the 25^th^ to 75^th^ percentile, whiskers span 10^th^ to 90^th^ percentile, and the line in the middle is the median. The individual points represent each measurement. Red and blue bands indicate the 25^th^ to 75^th^ percentile of maximum and minimum ratio values obtained for treatments with external H_2_O_2_ and DTT, to drive the sensor towards full oxidation and reduction. Data are from at least three independent experiments (Table S5). Letters denote statistical differences among treatments (Tukey’s Method, p-value < 0.05).

### H_2_S requires Ca^2+^ to induce stomatal closure

An additional hallmark of PTI signalling is a signature in cytosolic free calcium ([Ca^2+^]_cyt_) which triggers RBOH-activation directly by EF-hand binding and via Ca^2+^-dependent kinase signalling (Köster et al., 2022). Ca^2+^ signalling also contributes to other branches of the signalling network and shapes transcriptional re-programming. In guard cells, elicitors, like flg22, induce [Ca^2+^]_cyt_ oscillations, which are required for stomatal closure (Dodd et al., 2010; Thor and Peiter, 2014; Arnaud and Hwang, 2015; Thor et al., 2020). To address the question of whether H_2_S affects the Ca^2+^-branch of the stomatal immunity signalling network, we treated Col-0 epidermal peels with the H_2_S donor, GYY4137, in presence or absence of the membrane-permeable Ca^2+^chelator BAPTA-AM used to buffer cytosolic Ca^2+^ to suppress Ca^2+^ signaling, or the extracellular Ca^2+^ chelator, EGTA, to prevent any Ca^2+^ influx from the apoplast. Chelation of either intra- or extracellular Ca^2+^ prevented GYY4137 from inducing stomatal closure, suggesting that Ca^2+^ signalling is required for H_2_S induced stomatal closure (Figure 7).

**Figure 7:**
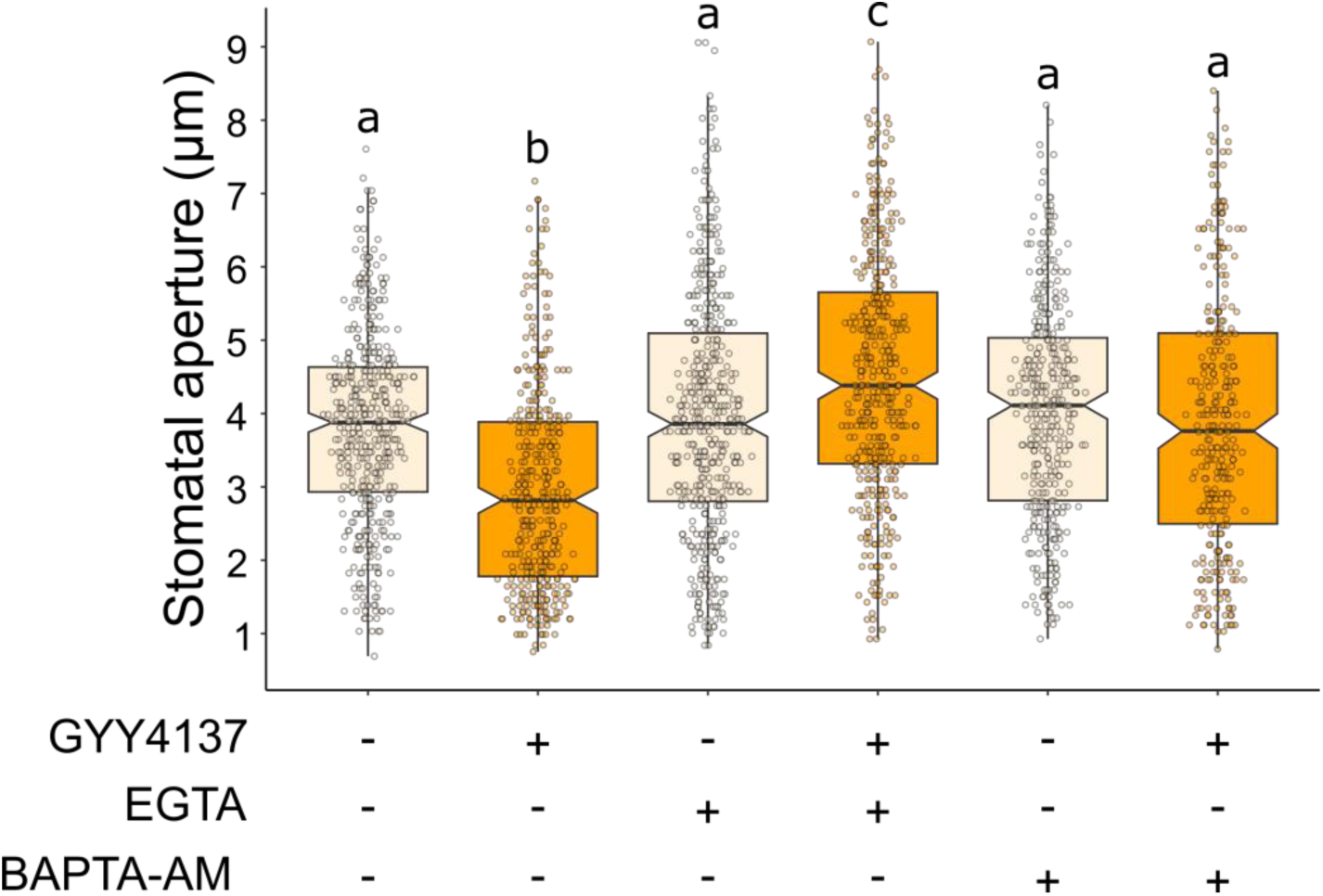
Apoplastic and cytosolic Ca^2+^ availability is required for H_2_S-induced stomatal closure. Epidermal peels from 4- to 5-week-old wild type (Col-0) Arabidopsis plants were pre-incubated in opening buffer (5 mM MES pH 6.1, 50 mM KCl) for 3 h under light and subsequently treated for 90 minutes with opening buffer, 25 µM of membrane permeable Ca^2+^ chelator, BAPTA-AM (BAPTA-AM), 200 µM of extracellular Ca^2+^ chelator EGTA (EGTA) in absence (-GYY4137) or presence (+ GYY4137) of 100 µM H_2_S donor, GYY4137 under light. The values of stomatal aperture are expressed in microns (µm) and represented in box-plots where the box is bound by the 25^th^ and 75^th^ percentile, whiskers span 10^th^ and 90^th^ percentile, and the line in the middle is the median. The individual points represent each measurement. Data are from at least three independent experiments (Table S5). Different letters denote statistical differences among treatments in each genotype (Tukey’s Method, p-value < 0.05).

## DISCUSSION

Stomatal closure is a key process of the plant immune response, since it generates a physical barrier that restricts pathogen entry to plant tissues. One of the first steps of this early response is the recognition of different PAMPs at the guard cell plasma membrane, an event that triggers PTI signalling involving several of the components that act as hubs within the guard cell signalling network (Hetherington and Woodward, 2003) and induces several *bona fide* defense processes like callose deposition (Zhang et al., 2020). In this study we demonstrated the involvement of H_2_S, and its main cytosolic source through DES1 activity, in the stomatal immunity response. We also present evidence that pinpoints where in the guard cell signalling network H_2_S may act.

### DES1 is required to induce stomatal closure under pathogen attack

H_2_S and different enzymatic sources, have been found to be involved in guard cell response to ABA, and other hormones related to abiotic stress (Liu and Xue, 2021; Pantaleno et al., 2021; Scuffi and García-Mata, 2021), while the role of H_2_S in the biotic stress response has been largely overlooked. Here, we use bacterial suspensions and PAMPs to show the involvement of DES1 in stomatal immunity. Mutant plants in *DES1* gene were impaired to close stomata in response to both bacteria and elicitors (flg22 and elf18). Moreover, stomata of *des1* closed when treated with flg22 together with H_2_S, supporting that DES1 participates in flg22-triggered response through the production of H_2_S (Figure 1 and 2). These results are in line with recent findings involving the mitochondrial source of H_2_S β-cyanoalanine synthase CAS-C1, where mutants lacking *CAS-C1* gene are unable to close stomata upon *Pst* and flg22 treatment (Pantaleno et al., 2024b). The involvement of DES1 in stomatal immunity suggests that *Pst* or flg22 might be modulating either the expression or the activity of DES1, as for ABA-dependent stomatal closure, where ABA induced *DES1* expression specifically in guard cells, and both DES1 and cytosolic H_2_S are required for stomatal closure (Scuffi et al., 2014; Zhang et al., 2019). In this case, however, there are not evident differences in *DES1* expression levels in control or flg22-treated guard cells (Figure S4), suggesting that flg22 might be regulating DES1 activity through a mechanism unknown until now. Recent reports on the mode of action of H_2_S in signal transduction processes indicate that H_2_S modulates target proteins through persulfidation. In this context, guard cells are not an exception, since the activity of several of the components that act as hubs in the guard cell signalling network were reported to be modulated by persulfidation (Reviewed in Pantaleno et al., 2021, Pantaleno and Scuffi., 2024). Such is also the case of the ABA-dependent response where DES1 is persulfidated and activated at the early stage of ABA treatment and can be reversibly and negatively regulated by H_2_O_2_ (Shen et al., 2020).

Given the relevance of stomatal movement dynamics for plant immunity, we investigated the *des1* response to *Pst* at the whole plant level by spray infection with *Pst* DC3000 *hrcC^-^* and we observed *des1* are less susceptible than Col-0 plants (Figure 3). This is in line with previous reports showing that plants lacking *DES1* presented a resistance phenotype to *Pst* DC3000 when leaf are infiltrated. The authors associate this response with the higher total glutathione and cysteine content in *des1* mutant, which can be linked with downstream immunity responses (Álvarez et al., 2012). In addition, a new line of evidences show that water availability in the apoplast can be crucial for pathogenesis (Aung et al., 2018). In consequence, it has been reported that when the stomata remain open, due to a lack of response to the closing stimulus, the apoplast water content which is necessary for sustain bacterial growth, is limited, generating phenotypes that are more resistant to pathogen’s attack (Freeman and Beattie, 2009; Xin et al., 2016; Liu et al., 2022).

### H_2_S and H_2_O_2_ involvement in stomatal immunity

The immune response triggered by flg22 is well characterized, not only in terms of the actors that compose it, but also in the temporal sequence of the processes involved. From this characterization, it is known that flg22 rapidly triggers an apoplastic ROS burst, highly dependent on the activation of RBOHD (Felix et al., 1999; Mersmann et al., 2010; Macho et al., 2012), which also has an active role in H_2_S and ABA-induced stomatal closure (Kwak et al., 2003; Scuffi et al., 2018; Shen et al., 2020). In fact, RBOHD was found to be persulfidated at Cys825 and Cys890 and activated, upon ABA treatment and these residues are required for ABA and H_2_S-dependent stomatal closure (Shen et al., 2020). Here we show that DES1 is partially required to induce apoplastic H_2_O_2_ in response to flg22. It would be interesting to measure apoplastic H_2_O_2_ production in response to exogenous application of H_2_S. Such an analysis is complicated given the inhibitory effect of H_2_S over the horseradish peroxidase (HRP) that is required for the luminol assay.

After the fast and transient apoplastic ROS burst, H_2_O_2_ is proposed to permeate into the cells through aquaporins, to amplify the defense response. Whether or not the

RBOHD-produced apoplastic ROS are indeed the exactly same molecules that cause cytosolic oxidation, is still a matter of debate and deserves future dissection. However, there is agreement in that both are required to induce stomatal closure (Kadota et al., 2014b; Toum et al., 2016; Arnaud et al., 2017; Rodrigues et al., 2017; Nietzel et al., 2019; Yang et al., 2021; Arnaud et al., 2023b). Here, we used the H_2_O_2_ biosensor, roGFP2-Orp1, and observed that flg22 induces cytosolic H_2_O_2_ increase with a time offset of about 50 min as compared to the apoplastic ROS burst. The cytosolic oxidation requires endogenous H_2_S and is partially dependent on DES1. Treatment with a general H_2_S donor also induces cytosolic H_2_O_2_ in the first 15 min in an RBOHD-independent manner (Figure 6) suggesting the involvement of an alternative source of cytosolic H_2_O_2_. These findings align with two recent studies; one demonstrating PAMP-mediated oxidation of roGFP2-Orp1 in the cytosol, independent of NADPH oxidases and apoplastic peroxidases (PRX), in Arabidopsis leaves (Arnaud et al., 2023a); and another one showing that roGFP2-Orp1 is oxidized by flg22 in Arabidopsis guard cells, even in *rbohD* and *rbohF* mutants (Arnaud et al., 2023b). In the latter study, RBOHF-dependent ROS release in guard cells was detected using the fluorescent dye H_2_DCFDA. This may suggest that ROS detected by H_2_DCFDA in undefined subcellular locations, but not by the cytosolic H_2_O_2_ biosensor, may contribute to flg22-dependent cytosolic ROS signalling (Arnaud et al., 2023a; Arnaud et al., 2023b). Similarly, PRX, rather than RBOHD or RBOHF, have been implicated in stomatal closure induced by cytokinin, and certain PRXs are strongly expressed in guard cells after flg22 treatment (Arnaud et al., 2017). Furthermore, PAMP-INDUCED PEPTIDE 1 (PIP1)-mediated stomatal closure is PRX-dependent (Hou et al., 2019). Weather PRXs, RBOHs, or other enzymes located in distinct subcellular compartments are involved in H_2_S signalling, and how their activity relates to stomatal immunity, remains to be elucidated. However, when *rbohD* plants are treated with the mitochondria-tagged-H_2_S donor AP39, a lower amount of cytosolic H_2_O_2_ is observed, suggesting that RBOHD is required to induce cytosolic H_2_O_2_ levels in response to mitochondrial H_2_S (Pantaleno et al., 2024b). The observed differences in H_2_O_2_ dynamics to a general or mitochondrial donor, may be attributed to the fact that H_2_S is a very reactive molecule and therefore, it is particularly likely to act close to its site of release (Filipovic et al., 2018; Benchoam et al., 2019). Furthermore, our findings suggest that different endogenous sources of H_2_S may operate at different levels to affect the stomatal response, highlighting the importance of the subcellular location in H_2_S-signalling.

### H_2_S participation in other PTI responses

Ca^2+^ is regarded as one of the few hubs in stomatal signaling network. Cytoplasmic Ca^2+^ increase is regulated by Ca^2+^ permeable channels, among them, CYCLIC NUCLEOTIDE GATED CHANNEL (CNGC) 2 and CNGC4, which assemble a channel that is phosphorylated upon flg22 perception, and the REDUCED HYPEROSMOLALITY INDUCED [Ca^2+^]_i_ INCREASE (OSCA) 1.3 which is also phosphorylated and required for pathogen-induced stomatal closure (Tian et al., 2019; Thor et al., 2020). The regulation of [Ca^2+^]_cyt_ modulates the activity of downstream targets containing Ca^2+^-binding domains, including calcium dependent protein kinases (CDPKs), NADPH oxidases and PHOSPHOLIPASES D (PLD). Moreover, once activated, CDPKs phosphorylate key targets as RBOHD and slow-type anion channel SLAC1 to induce stomatal closure (Qin and Wang, 2002; Ogasawara et al., 2008; Boudsocq et al., 2010; Geiger et al., 2010; Dubiella et al., 2013; Guzel Deger et al., 2015).

It is known that the second messengers H_2_O_2_ and Ca^2+^ have a complex interplay involving crossed regulation, where RBOHD-mediated apoplastic H_2_O_2_ production depends on flg22-triggered [Ca^2+^]_cyt_ increase, while Ca^2+^ entrance depends on apoplastic ROS burst (Marcec et al., 2019). Here we show that H_2_S is not able to close stomata when epidermal peels are pretreated with Ca^2+^ chelating agents (Figure 7) indicating that Ca^2+^ is required for H_2_S-induced stomatal closure. In ABA-dependent signalling it has been demonstrated that H_2_S activates S-type anion currents in a [Ca^2+^]_cyt_ and OST1-dependent manner, while OST1 persulfidation is necessary for its activation and for the Ca^2+^ influx from apoplast to the cytosol (Wang et al., 2016; Chen et al., 2020; Chen et al., 2021). OST1, together with anion channels SLAC1 and SLAC1 homolog 3 (SLAH3), are also needed to induce flg22-induced stomatal closure (Guzel Deger et al., 2015). Moreover, H_2_S inhibits inward-rectifying K^+^ channels in a Ca^2+^-independent way in tobacco guard cells (Papanatsiou et al., 2015). Interestingly, we show that DES1/H_2_S participate in these two processes that are core components of stomatal immunity response. Further studies will be needed to understand the mechanism by which H_2_S is acting upstream of Ca^2+^.

Another well characterized process acting downstream of flg22 signalling is the activation of MAPK and transcriptional reprogramming as a part of the PTI response (He et al., 2006; Boudsocq et al., 2010; Macho et al., 2012). Through guard cell specific-gene expression analysis and MAPK activity assay in seedlings, we show that DES1 is not involved in this pathway (Figure S3).

Through this work, we incorporate H_2_S into the mechanistic framework underlying plant defense response. We provide evidence for the involvement of DES1/H_2_S in stomatal immunity triggered by *Pst* and flg22. Our findings further elucidate the pathways within the stomatal signalling network that may be modulated by H_2_S.

In summary, we propose a model where the bacterial elicitor flg22 binds to its specific receptor FLS2, promoting the formation of FLS2-BAK1 complex, which triggers downstream signalling pathways, including the activation MAPK cascade. In parallel, DES1 is activated in the cytosol, catalyzing the production of ammonia, pyruvate, and, H_2_S from L-cysteine. DES1 is partially required to activate RBOHD via an unknown mechanism. RBOHD generates superoxide which is subsequently dismutated into H_2_O_2_ in the apoplast and transported into the cytosol via aquaporins. Additionally, H_2_S contributes to cytosolic H_2_O_2_ production through an undetermined source. Simultaneously, Ca^2+^ channels in the plasma membrane are activated, enabling Ca^2+^ influx into the cytosol to facilitate stomatal closure (Figure 8).

**Figure 8:**
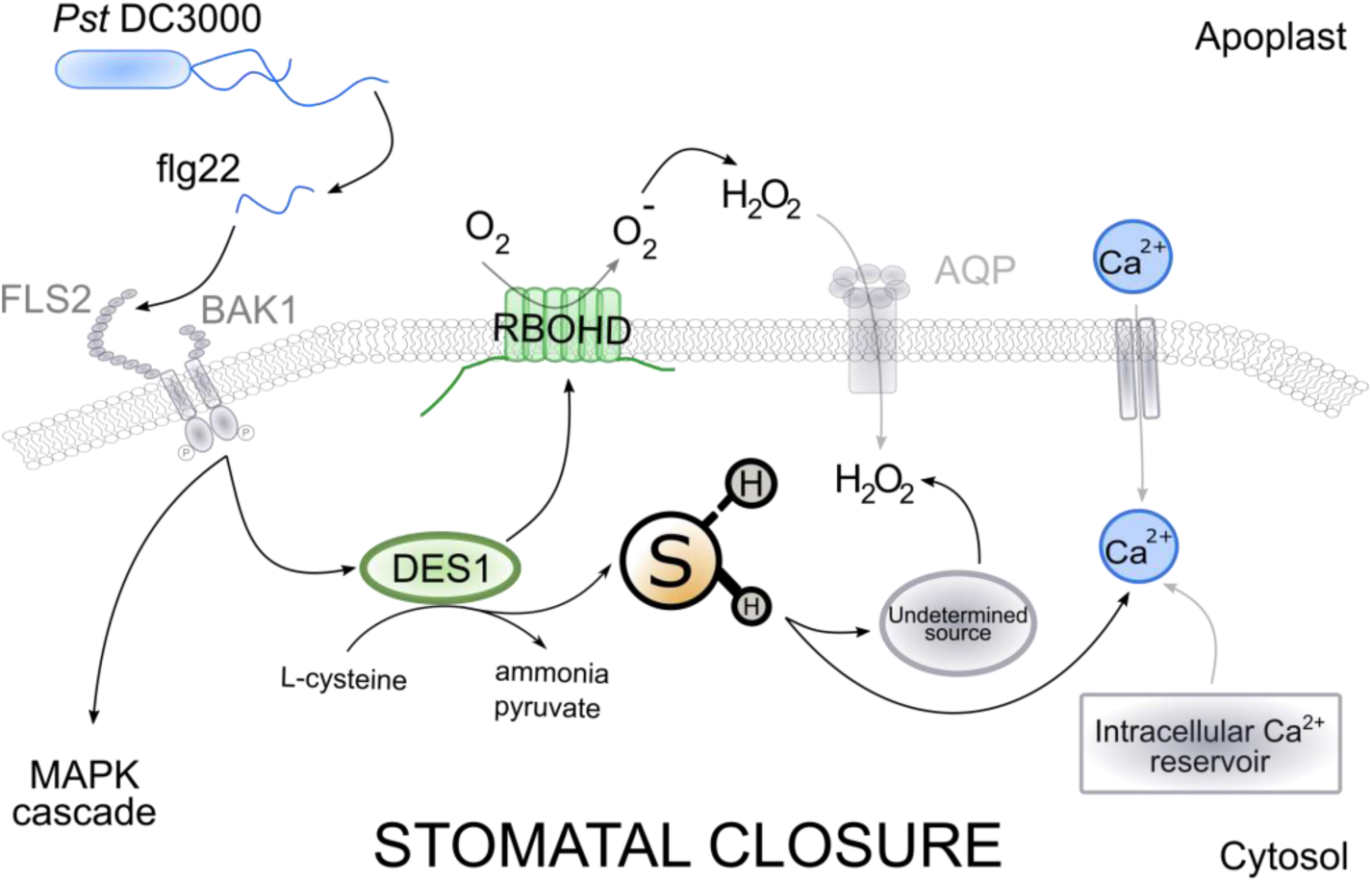
Working model of H_2_S and DES1-involvement in flg22-signalling in guard cells. Flagellin elicitor (flg22) from *Pseudomonas syringae* pv *tomato* DC3000 (*Pst* DC3000) binds to the specific receptor FLAGELLIN SENSING 2 (FLS2) and induce the formation of complex FLS2 and co-receptor BRI1-ASSOCIATED RECEPTOR KINASE 1 (BAK1) to triggers the signalling pathway which includes the activation of mitogen activated protein kinase (MAPK) cascade. On the other hand, upon flg22 perception *L*-CYSTEINE DESULFHIDRASE 1 (DES1) is activated generating ammonia, pyruvate and H_2_S from L-cysteine. DES1 is partially required to activate NADPH OXIDASE RESPIRATORY BURST OXIDASE HOMOLOG D (RBOHD) via an unknown mechanism. RBOHD produces superoxide (O_2_^-^) which is dismutated into H_2_O_2_ in the apoplast and then entry into the cytosol through aquaporins (AQP). Furthermore, H_2_S induces cytosolic H_2_O_2_ via an undetermined source. Ca^2+^ channels in the plasma membrane allowing Ca^2+^ to enter the cytosol to facilitate stomatal closure.

As such this study opens up future research avenues to identify the specific molecular targets for H_2_S and how they are modulated to integrate the stomatal immunity response.

## MATERIALS AND METHODS

### Biologic Material and Chemicals

*Arabidopsis thaliana* Columbia-0 (Col-0) wild type, *des1-1* (SALK_103855), *rbohD* (D3) (Torres et al., 2002) mutant and Col-0 expressing the roGFP2-Orp1 biosensor in the cytosol were available in our lab (Scuffi et al., 2018). *rbohD* and *des1* plants were crossed with the roGFP2-Orp1 biosensor lines. The F2 generation plants used was selected by fluorescence to ensure expression of the roGFP2-Orp1 sensor and by PCR-based genotyping to ensure homozygosity for the *rbohD* and *des1* locus (Primers listed in Table S5).

Seeds were germinated in soil (soil:vermiculite:perlite, 3:1:1) and kept at 4°C for 2 d. For stomatal aperture, apoplastic H_2_O_2_ detection, *Pst* spray-inoculation and gene expression experiments, plants were grown at 22°C in an 8-h-light/16-h-dark photoperiod at 200 µmol photons m^-2^ s^-1^ (IIB-CONICET-UNMdP. Mar del Plata, Argentina).

For biosensor analysis by epifluorescence microscopy, plants were grown at 22°C using an 8-h-light/16-h-dark photoperiod at 100 µmol photons m^-2^ s^-1^ (UNIMI, Milan, Italy).

For biosensor analysis by confocal microscopy, plants were grown at 25°C using a 16-h-light/8-h-dark photoperiod at 100 µmol photons m^-2^ s^-1^ (IBBP, Münster, Germany).

*Pseudomonas syringae* pv *tomato* DC3000 (*Pst* DC3000) and *Pseudomonas syringae* pv *tomato* DC3000 *hrcC^-^* (*Pst* DC3000 *hrcC*^-^) were kindly provided by Dr. Georgina Fabbro from CIQUIBIC-CONICET-UNC, Argentina. Bacteria were grown in King’s B media (2% (w/v) proteose peptone, 1% glycerol, 0.15% (w/v) K_2_HPO4, pH 7.2) supplemented with 100 mg L^-1^ rifampicin (for *Pst* DC3000 *hrcC^-^*) or 50 mg L^-1^ rifampicin and 50 mg L^-1^ kanamycin (for *Pst* DC3000) at 28°C.

The elicitors peptides flg22 (QRLSTGSRINSAKDDAAGLQIA) and elf18 (SKEKFERTKPHVNVGTIG) were purchased from ProteoGenix (Schiltigheim, France). The H_2_S donor, GYY4137 (morpholin-4-ium4 methoxyphenyl(morpholino) phosphinodithioate), the H_2_S scavenger, hypotaurine (HT), the horseradish peroxidase type VI-A (HRP), 3-aminophthalhydrazide, 5-amino-2,3-dihydro-1,4-phthalazinedione (luminol), MES buffer, EGTA and BAPTA-AM were purchased from Sigma-Aldrich.

For gene expression analysis, primers were synthesized by Macrogen (Seoul, Republic of Korea) and the fast Universal SYBR Green Master mix used was from Roche (Merck, Darmstadt, Alemania).

### Stomatal aperture assay

Stomatal aperture assays were performed according to Pantaleno et al. (2024a). Briefly, epidermal peels from abaxial side of fully expanded leaves from 5 to 6-week-old Arabidopsis plants were excised with tweezers and immediately floating in opening buffer (5 mM MES pH 6.1, 50 mM KCl) for 3 h under light (200 µmol photons m^-2^ s^-1^) and subsequently maintained in the same buffer omitting the active component (‘mock’) or exposed to different treatments as indicated in the legends. *Pst* DC3000 and *Pst* DC3000 *hrcC^-^* strains were grown at saturation at 28°C for 24 h in King’s B-agar plate and resuspended in sterile 10 mM MgCl_2_ until OD=0.1. Stomata were photographed using an AmScope MU1000 camera coupled to an Olympus CKX53 microscope with a 40x lens (LUCPlanFLN, 0.6 numerical aperture). The stomatal aperture width was quantified using the ImageJ analysis software (NIH, Bethesda, MD, USA)

### Bacterial Spray Inoculation

Five to 6-week-old Col-0 and *des1* plants were sprayed with a *Pst* DC3000 *hrcC^-^* suspension (OD = 0.2 [λ = 600 nm]) in 10 mM MgCl_2_, 0.02% (v/v) Silwet. Following spray inoculations, plants were kept covered with a transparent lid at 20°C under short-day conditions (8-h-light/16-h-dark photoperiod, 200 μmol photons m^−2^ s^−1^). Leaf discs samples (4 per plant) were taken at 24 and 72 hours (hpi) and the number of colony-forming units (CFU) was determined after serial dilution and plating as described (Johansson et al., 2014).

### Apoplastic H_2_O_2_ detection

Leaf discs from 4- to 5-week-old plants were cut and floated on deionized water overnight and then passed to 96-wells white plates. H_2_O_2_ production was triggered with 100 nM flg22 applied together with 20 mM luminol and 0.02 mg/L^-1^ HRP. Luminescence was measured with a luminometer (Thermo Scientific Luminoskan Ascent Microplate). Each plate was measured over a period of 30 min with a 2 min interval.

### Epifluorescence Microscopy

Biosensor analyses by epifluorescence microscopy was performed according to Pantaleno et al. 2024a. Briefly, epidermal peels from fully expanded leaf of Col-0 Arabidopsis 4- to 5-week-old plants expressing H_2_O_2_ biosensor roGFP2-Orp1 in the cytosol were floated in opening buffer for at list 7 h under light for recovery as we previously described (Scuffi et al., 2018). Peels were then treated with 1 µM flg22 preincubated or not for 10 min with 200 µM Hypotaurine (HT, H_2_S scavenger) for 1 h and to determine the dynamic range of the response of the roGFP2-based biosensors *in situ*, we used treatments with 10 mM H_2_O_2_ and 20 mM DTT for 10 min to induce full sensor oxidation and reduction, respectively. The peels were analyzed *in vivo* using an inverted Nikon Ti-E fluorescence microscope coupled to a Hamamatsu ORCA-D2 Dual CCD camera. Excitation light was produced by a fluorescent lamp Prior Lumen 200 PRO (Prior Scientific) and samples were imaged using a 60x oil immersion objective (CFI Plan APO Lambda 60x 1.4 numerical aperture). roGFP2-Orp1 was excited sequentially with 470/40 nm and 405/40 nm and the emission was collected using a 505/530 nm bandpass filter (GFP-specific filter) for both excitation wavelengths with a 2 x 2-pixel binning. The ratio 405/470 nm was calculated for each guard cell using the Ratio Plus plugin (https://imagej.net/ij/plugins/ratio-plus.html) for ImageJ analysis software (NIH, Bethesda, MD, USA). The background was subtracted for each channel and then the ROI was delimited to guard cells. Fluorescence was measured as the mean pixel intensity.

### Confocal Laser Scanning Microscopy

Biosensor analyses by CLSM was performed according to Pantaleno et al. 2024a. Briefly, epidermal peels from fully expanded leaf of Col-0, *des1* or *rbohD* from 4- to 5-week-old Arabidopsis plants expressing H_2_O_2_ biosensor roGFP2-Orp1 in the cytosol were floated in opening buffer for at least 7 h under light for recovery and then treated with 1 µM of flg22 or 100 µM of the H_2_S-donor GYY4137 according to the legend of the figure. To determine the dynamic range of the response of the roGFP2-based biosensors *in situ*, we used treatments with 10 mM H_2_O_2_ and 20 mM DTT to induce full sensor oxidation and reduction, respectively. The epidermal peels were mounted under a LSM980 inverted microscope (Carl Zeiss Microscopy). Images were collected with a 40x water immersion (C-Apochromat, 1.2 numerical aperture) and the biosensors were excited sequentially at 405 and 488 nm (line-switching mode) and emission was detected at 508 to 526 nm. Ratiometric images were analyzed using the custom MatLab program package, Redox Ratio Analysis (Fricker, 2016) (for *rbohD*) or ImageJ (NIH, Bethesda, MD, USA) (for *des1*) and ROI was set to specifically cover the guard cell.

### Gene expression in guard cell-enriched samples (GC-e)

Gene expression in GC-e was performed according to Pantaleno et al. 2024a. Briefly, Col-0 and *des1* Arabidopsis epidermal peels (GC-e, guard cell enriched) were floated in opening buffer (5 mM MES pH 6.1, 50 mM KCl) for 3 h under light and then treated with 1 µM flg22 for 60 min in the same conditions as we previously described (Scuffi et al., 2014). Total RNA was extracted using homemade reagent: Phenol saturated in buffer Tris pH 8 38 % v/v, Guanidine thiocyanate 11.8 % w/v, Ammonium thicyanate 7.91 % w/v, Sodium acetate 0.1 M pH 5, Glycerol 5% v/v. Subsequently, 1 µg of total RNA was used for the RT-qPCR reaction using an oligo(dT) primer and Moloney murine leukemia virus reverse transcriptase (M-MLV RT, SIGMA).

For qPCR reaction, Fast Universal SYBR Green Master mix was employed, using a Step-One Real Time PCR machine from Applied Biosystems. The standard amplification program was used. The expression levels of the gene of interest were normalized to those of the constitutive *ACT2* (*At3g18780*) gene by subtracting the cycle threshold value of *ACT2* from the cycle threshold value of the gene (ΔCT). The nucleotide sequences of the specific primers for qPCR analysis are listed in Supplemental Table S5. The annealing temperature for each primer was 60°C. LinRegPCR was the program employed for the analysis of RT-qPCR data (Ruijter et al., 2009).

### MAPK Activation

MAPK assay was performed on nine 2-week-old Arabidopsis Col-0 and *des1* seedlings grown in liquid MS plus 1% (w/v) sucrose at 25°C using a 16-h-light/8-h-dark photoperiod at 100 µmol photons m^-2^ s^-1^. Seedlings were treated with 1 µM flg22 for 0, 5, 15, or 30 min and flash-frozen in liquid nitrogen. MAPK activation was monitored by western blotting with antibodies that recognize the dual phosphorylation of the activation loop of MAPK (pTEpY). Phospho-p44/42 MAPK (Erk1/2; Thr-202/Tyr-204) rabbit monoclonal antibodies (Cell Signalling) were used according to the manufacturer’s protocol (1:5,000). Blots were stained with Ponceau S to verify equal loading.

### Statistical analysis

Data analyses were performed using RStudio (R Foundation for Statistical Computing, Vienna, Austria; URL: https://www.Rproject.org/). The statistically significant differences were analyzed using Student’s t-test, Generalized Linear Model or Generalized Linear Mix Model procedure, with the gls function from the nlme library. Multiple comparisons among individual means were performed by Tukey’s Method. The error distribution was Gaussian or Gamma and all effect were considered significative at p < 0.05 or p < 0.001 (Table S5).

## Acknowledgements

We thank the Deutsche Forschungsgemeinschaft (DFG) for funding through the infrastructure grant <u>INST 211/903-1 FUGG</u> for the confocal microscope as operated by the Imaging Network of the University of Münster (RI_00497), the University of Mar del Plata (UNMdP), Consejo Nacional de Investigaciones Científicas y Técnicas (CONICET), the Agencia Nacional de Promoción Científica y Tecnológica (ANPCyT; PICT 2016 N° 2553, PICT 2017 N° 601, 2018 N° 1449, PICT 2019 N° 1040 and PICT 2021 N° 92), the DAAD for Research Stays for University Academics and Scientists, 2020 (57507437) fellowship to D.S. and Travelling Fellowship from The Company of Biologists to R.P. Part of the work was carried out with the support of the NOLIMITS Center of Excellence for Plant Biology and Other Life Sciences established by the University of Milan.

## Author Contributions

DS performed and designed most of the experiment, analyzed the data and wrote the article, RP performed the experiments from figure 3 and S2, PS performed the experiments from figure 3 and 7, J-ON performed the experiments from figure 5, AC, MS and AL participate in the experimental design, discussion and writing and CGM conceived the project, analyzed data and wrote the article. All authors contribute to the writing of the article

## Bibliography

Agurla S, Gayatri G, Raghavendra AS (2014) Nitric oxide as a secondary messenger during stomatal closure as a part of plant immunity response against pathogens. Nitric Oxide 43: 89–96

Álvarez C, Ángeles Bermúdez M, Romero LC, Gotor C, García I (2012) Cysteine homeostasis plays an essential role in plant immunity. New Phytologist 193: 165– 177

Álvarez C, Calo L, Romero LC, García I, Gotor C (2010) An O - Acetylserine(thiol)lyase Homolog with l -Cysteine Desulfhydrase Activity Regulates Cysteine Homeostasis in Arabidopsis. Plant Physiology 152: 656–669

Arnaud D, Deeks MJ, Smirnoff N (2023a) Organelle-targeted biosensors reveal distinct oxidative events during pattern-triggered immune responses. Plant Physiology 191: 2551–2569

Arnaud D, Deeks MJ, Smirnoff N (2023b) RBOHF activates stomatal immunity by modulating both reactive oxygen species and apoplastic pH dynamics in Arabidopsis. Plant Journal 1–12

Arnaud D, Hwang I (2015) A sophisticated network of signaling pathways regulates stomatal defenses to bacterial pathogens. Molecular Plant 8: 566–581

Arnaud D, Lee S, Takebayashi Y, Choi D, Choi J, Sakakibara H, Hwang I (2017) Cytokinin-mediated regulation of reactive oxygen species homeostasis modulates stomatal immunity in arabidopsis. Plant Cell 29: 543–559

Aroca A, Benito JM, Gotor C, Romero LC (2017) Persulfidation proteome reveals the regulation of protein function by hydrogen sulfide in diverse biological processes in Arabidopsis. Journal of Experimental Botany 68: 4915–4927

Aroca Á, Serna A, Gotor C, Romero LC (2015) S-Sulfhydration: A Cysteine Posttranslational Modification in Plant Systems. Plant Physiology 168: 334–342

Aroca A, Zhang J, Xie Y, Romero LC, Gotor C (2021) Hydrogen sulfide signaling in plant adaptations to adverse conditions: molecular mechanisms. Journal of Experimental Botany 72: 5893–5904

Aung K, Jiang Y, He SY (2018) The role of water in plant–microbe interactions. The Plant Journal 93: 771–780

Benchoam D, Cuevasanta E, Möller M, Alvarez B (2019) Hydrogen Sulfide and Persulfides Oxidation by Biologically Relevant Oxidizing Species. Antioxidants 8: 48

Bethke G, Pecher P, Eschen-Lippold L, Tsuda K, Katagiri F, Glazebrook J, Scheel D, Lee J (2012) Activation of the Arabidopsis thaliana Mitogen-Activated Protein Kinase MPK11 by the Flagellin-Derived Elicitor Peptide, flg22. Molecular Plant-Microbe Interactions 25: 471–480

Bjornson M, Pimprikar P, Nürnberger T, Zipfel C (2021) The transcriptional landscape of Arabidopsis thaliana pattern-triggered immunity. Nature Plants 7: 579–586

Blatt MR (2000) Cellular signaling and volume control in stomatal movements in plants. Annual review of cell and developmental biology 16: 221–41

Bloem E (2004) Sulphur supply and infection with Pyrenopeziza brassicae influence L-cysteine desulphydrase activity in Brassica napus L. Journal of Experimental Botany 55: 2305–2312

Bloem E, Haneklaus S, Kesselmeier J, Schnug E (2012) Sulfur fertilization and fungal infections affect the exchange of H(2)S and COS from agricultural crops. Journal of agricultural and food chemistry 60: 7588–96

Bloem E, Haneklaus S, Salac I, Wickenhäuser P, Schnug E (2007) Facts and fiction about sulfur metabolism in relation to plant-pathogen interactions. Plant Biology 9: 596–607

Boudsocq M, Willmann MR, McCormack M, Lee H, Shan L, He P, Bush J, Cheng S-H, Sheen J (2010) Differential innate immune signalling via Ca2+ sensor protein kinases. Nature 464: 418–422

Chen S, Jia H, Wang X, Shi C, Wang X, Ma P, Wang J, Ren M, Li J (2020) Hydrogen Sulfide Positively Regulates Abscisic Acid Signaling through Persulfidation of SnRK2.6 in Guard Cells. Molecular Plant 13: 732–744

Chen S, Wang X, Jia H, Li F, Ma Y, Liesche J, Liao M, Ding X, Liu C, Chen Y, et al (2021) Persulfidation-induced structural change in SnRK2.6 establishes intramolecular interaction between phosphorylation and persulfidation. Molecular Plant 14: 1814–1830

Daudi A, Cheng Z, O’Brien JA, Mammarella N, Khan S, Ausubel FM, Paul Bolwell G (2012) The apoplastic oxidative burst peroxidase in Arabidopsis is a major component of pattern-triggered immunity. Plant Cell 24: 275–287

Dodd AN, Kudla J, Sanders D (2010) The language of calcium signaling. Annual Review of Plant Biology 61: 593–620

Du X, Jin Z, Zhang L, Liu X, Yang G, Pei Y (2019) H2S is involved in ABA-mediated stomatal movement through MPK4 to alleviate drought stress in Arabidopsis thaliana. Plant Soil 435: 295–307

Dubiella U, Seybold H, Durian G, Komander E, Lassig R, Witte C-P, Schulze WX, Romeis T (2013) Calcium-dependent protein kinase/NADPH oxidase activation circuit is required for rapid defense signal propagation. Proceedings of the National Academy of Sciences of the United States of America 110: 8744–9

Felix G, Duran JD, Volko S, Boller T (1999) Plants have a sensitive perception system for the most conserved domain of bacterial flagellin. The Plant Journal 18: 265–276

Filipovic MR, Zivanovic J, Alvarez B, Banerjee R (2018) Chemical Biology of H2S Signaling through Persulfidation. Chemical Reviews 118: 1253–1337

Freeman BC, Beattie GA (2009) Bacterial Growth Restriction During Host Resistance to *Pseudomonas syringae* Is Associated with Leaf Water Loss and Localized Cessation of Vascular Activity in *Arabidopsis thaliana*. MPMI 22: 857–867

Fricker MD (2016) Quantitative Redox Imaging Software. Antioxidants & redox signaling 24: 752–62

Geiger D, Scherzer S, Mumm P, Marten I, Ache P, Matschi S, Liese A, Wellmann C, Al-Rasheid KAS, Grill E, et al (2010) Guard cell anion channel SLAC1 is regulated by CDPK protein kinases with distinct Ca2+ affinities. Proceedings of the National Academy of Sciences of the United States of America 107: 8023–8

Guzel Deger A, Scherzer S, Nuhkat M, Kedzierska J, Kollist H, Brosché M, Unyayar S, Boudsocq M, Hedrich R, Roelfsema MRG (2015) Guard cell SLAC 1-type anion channels mediate flagellin-induced stomatal closure. New Phytologist 208: 162–173

Hauck P, Thilmony R, He SY (2003) A Pseudomonas syringae type III effector suppresses cell wall-based extracellular defense in susceptible Arabidopsis plants. Proceedings of the National Academy of Sciences 100: 8577–8582

He P, Shan L, Lin N-C, Martin GB, Kemmerling B, Nürnberger T, Sheen J (2006) Specific Bacterial Suppressors of MAMP Signaling Upstream of MAPKKK in Arabidopsis Innate Immunity. Cell 125: 563–575

Hetherington AM, Woodward FI (2003) The role of stomata in sensing and driving environmental change. Nature 424: 901–908

Hou S, Shen H, Shao H (2019) PAMP-induced peptide 1 cooperates with salicylic acid to regulate stomatal immunity in *Arabidopsis thaliana*. Plant Signaling & Behavior 14: 1666657

Kadota Y, Sklenar J, Derbyshire P, Stransfeld L, Asai S, Ntoukakis V, Jones JD, Shirasu K, Menke F, Jones A, et al (2014a) Direct Regulation of the NADPH Oxidase RBOHD by the PRR-Associated Kinase BIK1 during Plant Immunity. Molecular Cell 54: 43–55

Kadota Y, Sklenar J, Derbyshire P, Stransfeld L, Asai S, Ntoukakis V, Jones JD, Shirasu K, Menke F, Jones A, et al (2014b) Direct Regulation of the NADPH Oxidase RBOHD by the PRR-Associated Kinase BIK1 during Plant Immunity. Molecular Cell 54: 43–55

Kim T-H, Böhmer M, Hu H, Nishimura N, Schroeder JI (2010) Guard cell signal transduction network: advances in understanding abscisic acid, CO2, and Ca2+ signaling. Annual review of plant biology 61: 561–91

Köster P, DeFalco TA, Zipfel C (2022) Ca ^2+^ signals in plant immunity. The EMBO Journal 41: e110741

Kwak JM, Mori IC, Pei ZM, Leonhard N, Angel Torres M, Dangl JL, Bloom RE, Bodde S, Jones JDG, Schroeder JI (2003) NADPH oxidase AtrbohD and AtrbohF genes function in ROS-dependent ABA signaling in arabidopsis. EMBO Journal 22: 2623–2633

Li L, Li M, Yu L, Zhou Z, Liang X, Liu Z, Cai G, Gao L, Zhang X, Wang Y, et al (2014) The FLS2-Associated Kinase BIK1 Directly Phosphorylates the NADPH Oxidase RbohD to Control Plant Immunity. Cell Host & Microbe 15: 329–338

Liu H, Xue S (2021) Interplay between hydrogen sulfide and other signaling molecules in the regulation of guard cell signaling and abiotic/biotic stress response. Plant Communications 2: 100179

Liu Z, Hou S, Rodrigues O, Wang P, Luo D, Munemasa S, Lei J, Liu J, Ortiz-Morea FA, Wang X, et al (2022) Phytocytokine signalling reopens stomata in plant immunity and water loss. Nature 605: 332–339

Macho AP, Boutrot F, Rathjen JP, Zipfel C (2012) ASPARTATE OXIDASE plays an important role in Arabidopsis stomatal immunity. Plant Physiology 159: 1845– 1856

Marcec MJ, Gilroy S, Poovaiah BW, Tanaka K (2019) Mutual interplay of Ca2+ and ROS signaling in plant immune response. Plant Science 283: 343–354

Melotto M, Fochs B, Jaramillo Z, Rodrigues O (2024) Fighting for Survival at the Stomatal Gate. Annual Review of Plant Biology 75: 551–577

Melotto M, Underwood W, Koczan J, Nomura K, He SY (2006) Plant Stomata Function in Innate Immunity against Bacterial Invasion. Cell 126: 969–980

Melotto M, Zhang L, Oblessuc PR, He SY (2017) Stomatal Defense a Decade Later. Plant Physiology 174: 561–571

Mersmann S, Bourdais G, Rietz S, Robatzek S (2010) Ethylene signaling regulates accumulation of the FLS2 receptor and is required for the oxidative burst contributing to plant immunity. Plant physiology 154: 391–400

Nietzel T, Els M, Ruberti C, Steinbeck J, Moerschbacher BM, Costa A, Fricker MD, Stefanie JM, Meyer AJ, Schwarzl M (2019) Methods The fluorescent protein sensor roGFP2-Orp1 monitors in vivo H 2 O 2 and thiol redox integration and elucidates intracellular H 2 O 2 dynamics during elicitor-induced oxidative burst in Arabidopsis. 1649–1664

Ogasawara Y, Kaya H, Hiraoka G, Yumoto F, Kimura S, Kadota Y, Hishinuma H, Senzaki E, Yamagoe S, Nagata K, et al (2008) Synergistic Activation of the Arabidopsis NADPH Oxidase AtrbohD by Ca2+ and Phosphorylation. Journal of Biological Chemistry 283: 8885–8892

Pantaleno R, Schiel P, García-Mata C, Scuffi D (2024a) Analysis of Guard Cell Readouts Using Arabidopsis thaliana Isolated Epidermal Peels. BIO-PROTOCOL. doi: 10.21769/BioProtoc.5033

Pantaleno R, Scuffi D, García-Mata C (2021) Hydrogen sulphide as a guard cell network regulator. New Phytologist 230: 451–456

Pantaleno R, Scuffi D, Schiel P, Schwarzländer M, Costa A, García-Mata C (2024b) Mitochondrial ß-Cyanoalanine Synthase Participates in flg22-Induced Stomatal Immunity. Plant Cell & Environment pce.15155

Pantaleno R, Scuffi D (2024) Posttranslational modifications triggered by H2S in plant cells. *In* Modolo LV, Da-Silva CJ, eds, H2S in Plants: Past, Present and Beyond. Elsevier Inc, pp 169–191

Papanatsiou M, Scuffi D, Blatt MR, García-Mata C (2015) Hydrogen Sulfide Regulates Inward-Rectifying K + Channels in Conjunction with Stomatal Closure. Plant Physiology 168: 29–35

Qi J, Song C, Wang B, Zhou J, Kangasjärvi J, Zhu J, Gong Z (2018) Reactive oxygen species signaling and stomatal movement in plant responses to drought stress and pathogen attack. JIPB 60: 805–826

Qin C, Wang X (2002) The Arabidopsis phospholipase D family. Characterization of a calcium-independent and phosphatidylcholine-selective PLD zeta 1 with distinct regulatory domains. Plant physiology 128: 1057–1068

Rodrigues O, Reshetnyak G, Grondin A, Saijo Y, Leonhardt N, Maurel C, Verdoucq L (2017) Aquaporins facilitate hydrogen peroxide entry into guard cells to mediate ABA- and pathogen-triggered stomatal closure. Proceedings of the National Academy of Sciences of the United States of America 114: 9200–9205

Ruijter JM, Ramakers C, Hoogaars WMH, Karlen Y, Bakker O, van den Hoff MJB, Moorman a FM (2009) Amplification efficiency: linking baseline and bias in the analysis of quantitative PCR data. Nucleic acids research 37: e45

Schroeder JI, Allen GJ, Hugouvieux V, Kwak JM, Waner D (2001) GUARD CELL SIGNAL TRANSDUCTION. Annual review of plant physiology and plant molecular biology 52: 627–658

Scuffi D, Álvarez C, Laspina N, Gotor C, Lamattina L, García-Mata C (2014) Hydrogen Sulfide Generated by l-Cysteine Desulfhydrase Acts Upstream of Nitric Oxide to Modulate Abscisic Acid-Dependent Stomatal Closure. Plant Physiology 166: 2065–2076

Scuffi D, García-Mata C (2021) Hydrogen Sulfide and Stomatal Movement. *In* MN Khan, MH Siddiqui, S Alamri, FJ Corpas, eds, Hydrogen Sulfide and Plant Acclimation to Abiotic Stresses. Springer International Publishing, Cham, pp 87– 107

Scuffi D, Nietzel T, Di Fino LM, Meyer AJ, Lamattina L, Schwarzländer M, Laxalt AM, García-Mata C (2018) Hydrogen Sulfide Increases Production of NADPH Oxidase-Dependent Hydrogen Peroxide and Phospholipase D-Derived Phosphatidic Acid in Guard Cell Signaling. Plant Physiology 176: 2532–2542

Shen J, Zhang J, Zhou M, Zhou H, Cui B, Gotor C, Romero LC, Fu L, Yang J, Foyer CH, et al (2020) Persulfidation-based Modification of Cysteine Desulfhydrase and the NADPH Oxidase RBOHD Controls Guard Cell Abscisic Acid Signaling. Plant Cell 32: 1000–1017

Shi H, Ye T, Han N, Bian H, Liu X, Chan Z (2015) Hydrogen sulfide regulates abiotic stress tolerance and biotic stress resistance in Arabidopsis. Journal of Integrative Plant Biology 57: 628–640

Soto D, Córdoba JP, Villarreal F, Bartoli C, Schmitz J, Maurino VG, Braun HP, Pagnussat GC, Zabaleta E (2015) Functional characterization of mutants affected in the carbonic anhydrase domain of the respiratory complex I in Arabidopsis thaliana. The Plant Journal 83: 831–844

Thor K, Jiang S, Michard E, George J, Scherzer S, Huang S, Dindas J, Derbyshire P, Leitão N, DeFalco TA, et al (2020) The calcium-permeable channel OSCA1.3 regulates plant stomatal immunity. Nature 585: 569–573

Thor K, Peiter E (2014) Cytosolic calcium signals elicited by the pathogen-associated molecular pattern flg22 in stomatal guard cells are of an oscillatory nature. New Phytologist 204: 873–881

Tian W, Hou C, Ren Z, Wang C, Zhao F, Dahlbeck D, Hu S, Zhang L, Niu Q, Li L, et al (2019) A calmodulin-gated calcium channel links pathogen patterns to plant immunity. Nature. doi: 10.1038/s41586-019-1413-y

Torres MA, Dangl JL, Jones JDG (2002) Arabidopsis gp91phox homologues AtrbohD and AtrbohF are required for accumulation of reactive oxygen intermediates in the plant defense response. Proceedings of the National Academy of Sciences 99: 517–522

Toum L, Torres PS, Gallego SM, Benavídes MP, Vojnov AA, Gudesblat GE (2016) Coronatine Inhibits Stomatal Closure through Guard Cell-Specific Inhibition of NADPH Oxidase-Dependent ROS Production. Frontiers in Plant Science 7: 1–12

Underwood W, Melotto M, He SY (2007) Role of plant stomata in bacterial invasion. Cellular Microbiology 9: 1621–1629

Vojtovič D, Luhová L, Petřivalský M (2021) Something smells bad to plant pathogens: Production of hydrogen sulfide in plants and its role in plant defence responses. Journal of Advanced Research 27: 199–209

Wang L, Wan R, Shi Y, Xue S (2016) Hydrogen Sulfide Activates S-Type Anion Channel via OST1 and Ca 2+ Modules. Molecular Plant 9: 489–491

Wang P, Fang H, Gao R, Liao W (2021) Protein Persulfidation in Plants: Function and Mechanism. Antioxidants 10: 1631

Xin X-F, Nomura K, Aung K, Velásquez AC, Yao J, Boutrot F, Chang JH, Zipfel C, He SY (2016) Bacteria establish an aqueous living space in plants crucial for virulence. Nature 539: 524–529

Yang Q, Peng Z, Ma W, Zhang S, Hou S, Wei J, Dong S, Yu X, Song Y, Gao W, et al (2021) Melatonin functions in priming of stomatal immunity in Panax notoginseng and Arabidopsis thaliana. Plant Physiology 187: 2837–2851

Zhang J, Coaker G, Zhou JM, Dong X (2020) Plant Immune Mechanisms: From Reductionistic to Holistic Points of View. Molecular Plant 13: 1358–1378

Zhang J, Zhou M, Ge Z, Shen J, Zhou C, Gotor C, Romero LC, Duan X, Liu X, Wu D, et al (2019) ABA-triggered guard cell L -cysteine desulfhydrase function and in situ H 2 S production contributes to heme oxygenase-modulated stomatal closure . Plant, Cell & Environment 1–13

