## Supplemental material for "Hydrogen Sulfide modulates Flagellin-Induced Stomatal Immunity"

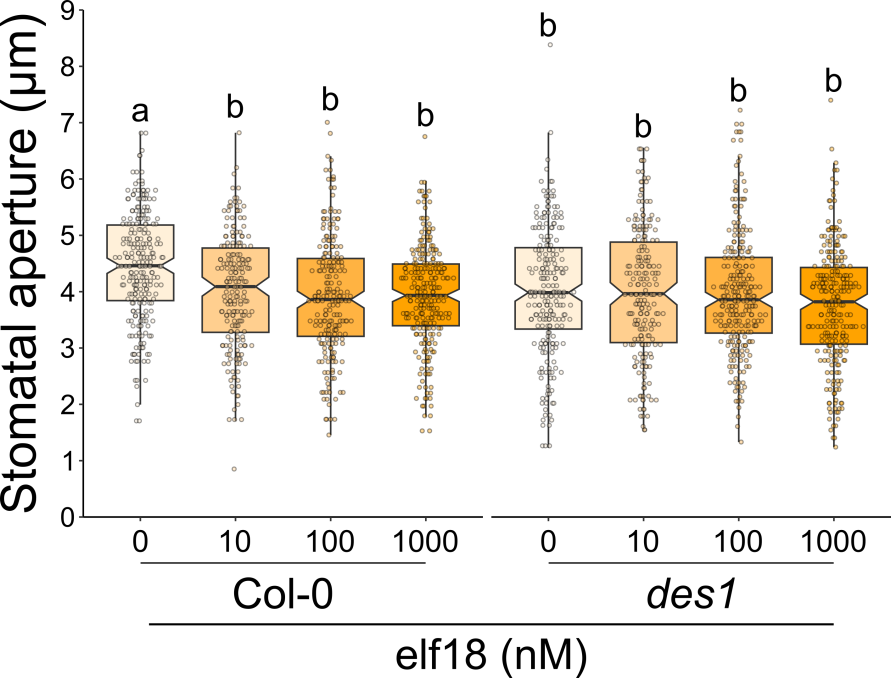

**Supplemental S1: DES1 is required for elf18-induced stomatal closure response.** Epidermal peels from wild type (Col-0) and *des1* (*des1*) mutant Arabidopsis plants were pre-incubated in opening buffer (5 mM MES pH 6.1, 50 mM KCl) for 3 h under light and subsequently treated for 90 minutes with 0, 10, 100 or 1000 nM elf18 in the same buffer under light. The values of stomatal aperture are expressed in microns (µm) and represented in box-plots where the box is bound by the 25^th^ and 75^th^ percentile, whiskers span 10^th^ and 90^th^ percentile, and the line in the middle is the median. The individual points represent each measurement. Data are from at least three independent experiments (Table S4). Different letters denote statistical differences among treatments (Tukey’s Method, p-value < 0.05)

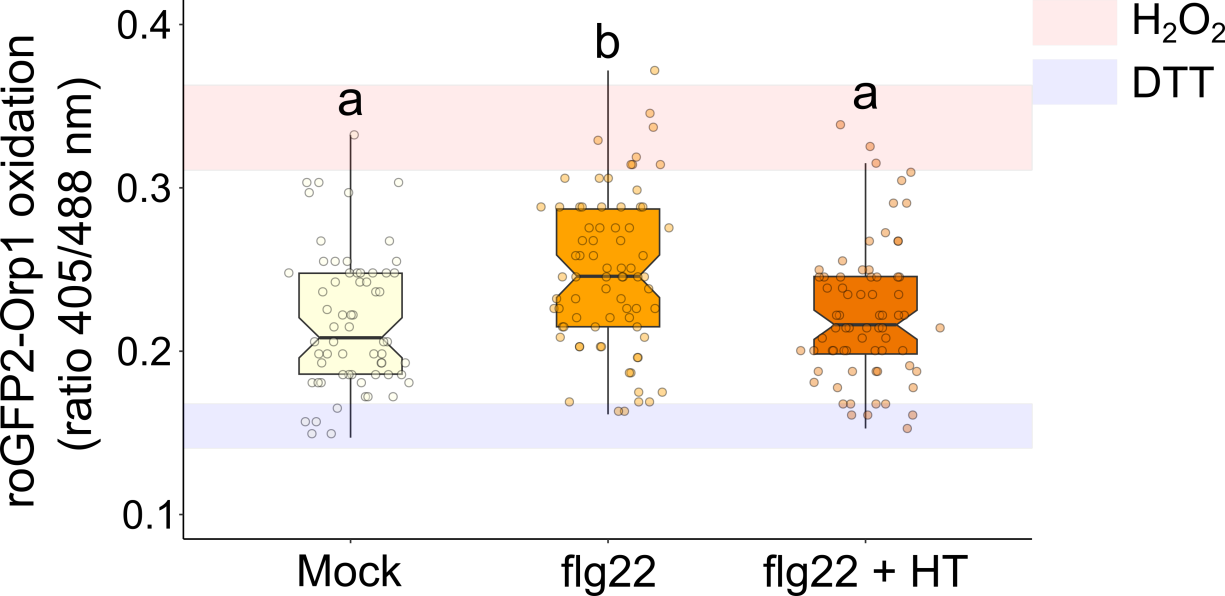

**Supplemental S2: Endogenous H_2_S is required for flg22-dependent cytosolic H_2_O_2_ production in guard cells.** Epidermal peels from 4- to 5-week-old Col-0 plants expressing H_2_O_2_ specific biosensor roGFP2-Orp1 in the cytosol were incubated for 7-12 h in opening buffer (5 mM MES pH 6.1, 50 mM KCl) and subsequently treated with opening buffer (Mock), 1 µM flg22 (flg22) for 60 min, or pre-incubated with 200 µM of H_2_S scavenger hypotaurine for 10 min and then treated with 1 µM flg22 (flg22 + HT). Moreover, epidermal peels were treated with 20 mM DTT or 10 mM H_2_O_2_ for 10 min to estimate the dynamic range of the sensor in situ. Values are expressed as the ratio of 405/488 nm and are represented in the box plots where the box is bound by the 25^th^ to 75^th^ percentile, whiskers span 10^th^ to 90^th^ percentile, and the line in the middle is the median. The individual points represent each measurement. Red and blue bands indicate the 25^th^ to 75^th^ percentile of maximum and minimum ratio values obtained for treatments with external H_2_O_2_ and DTT, to drive the sensor towards full oxidation and reduction. Data are from at least three independent experiments (Table S5). Different letters denote statistical differences among treatments (Tukey’s Method, p-value < 0.05)

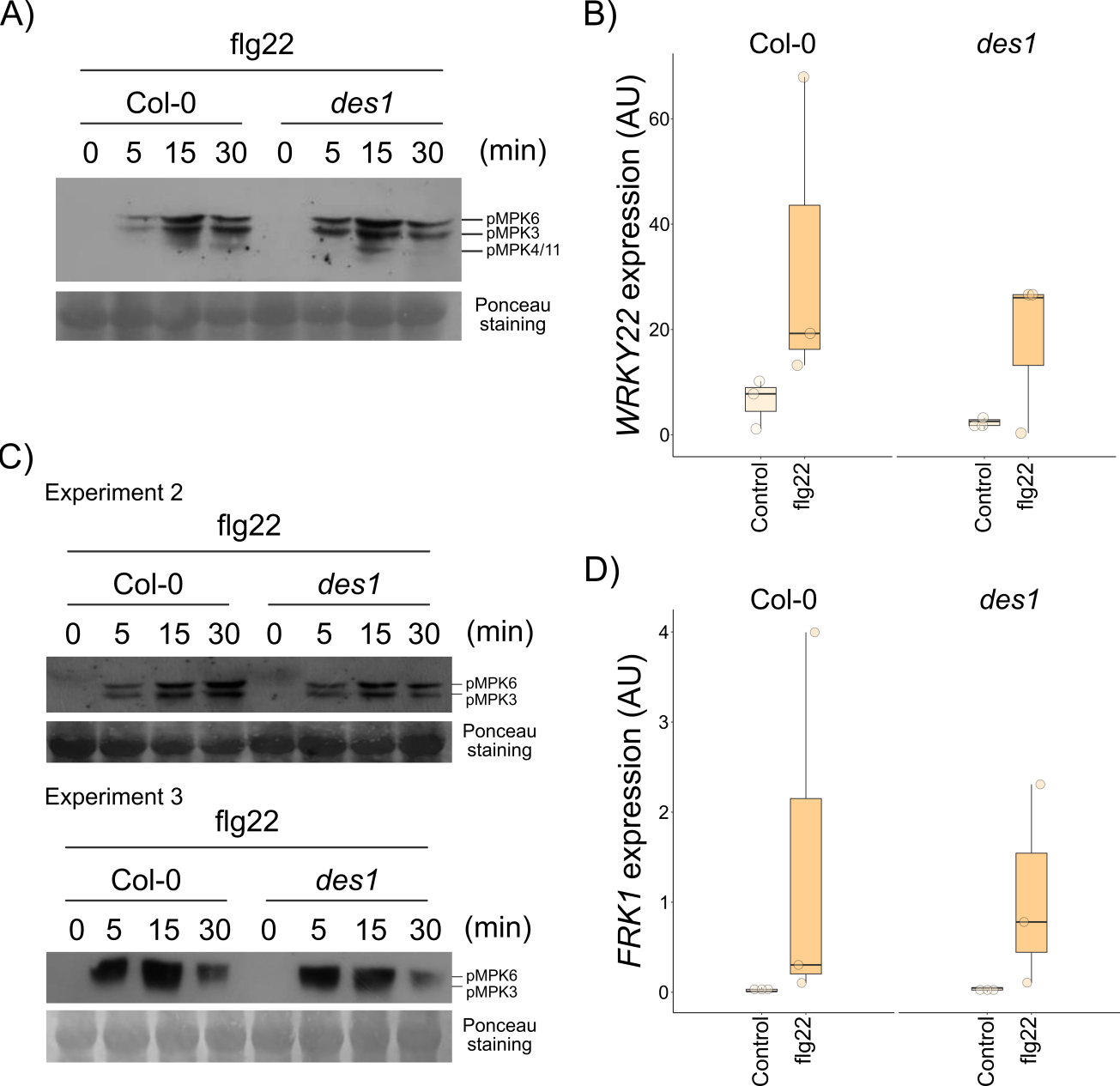

**Figure S3: DES1 is not involved in MAPKs activation.** Col-0 and *des1* 2-week-old seedlings were grown in liquid half-strength MS medium, treated with 1 µM flg22 for 0, 5, 15 and 30 minutes, and an immunological assay for detecting the activated mitogen-activated protein kinases (MAPKs) was performed. A representative immunoblot of three experiments is shown (A). The other two experimental replicates are shown in (C). Epidermal strips prepared from 4-6-week-old wild type (Col-0) and *des1* (*des1*) plants were preincubated for 3 h in opening buffer (5 mM MES, pH 6.1, 50 mM KCl) and then treated with (flg22) or without (Control) 1 µM flg22 under light. After 60 min of treatment, qRT-PCR analysis of *WRKY22* (B) and *FRK1* (D) genes expression was performed in guard cell enriched (GC-e) RNA samples. Values are expressed in arbitrary units (AU) and are represented in box-plots where the box is bound by the 25^th^ to 75^th^ percentile, whiskers span 10^th^ to 90^th^ percentile, and the line in the middle is the median. The individual points represent each measurement (Table S4).

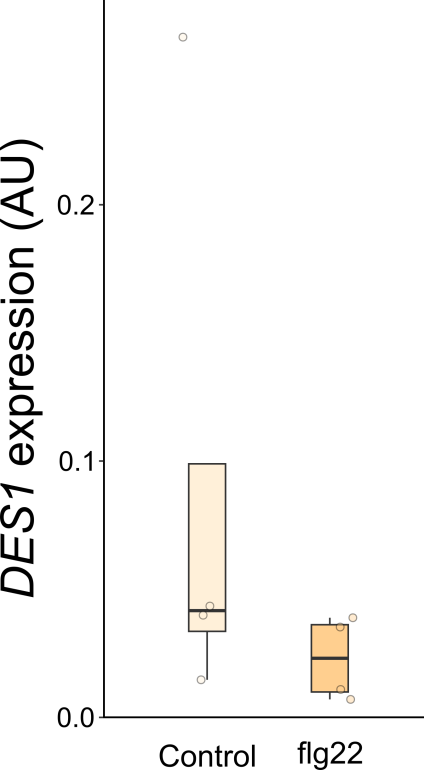

**Supplemental S4: *DES1* expression is not affected in GC-e samples under flg22 treatment.** Epidermal strips prepared from 4-6-week-old wild type (Col-0) plants were preincubated for 3 h in opening buffer (5 mM MES, pH 6.1, 50 mM KCl) and then treated with (flg22) or without (Control) 1 µM flg22 under light. After 60 min of treatment, RT-qPCR analysis of *DES1* gene expression was performed in GC-e RNA. Values are expressed in arbitrary units (AU) and are represented in box-plots where the box is bound by the 25^th^ to 75^th^ percentile, whiskers span 10^th^ to 90^th^ percentile, and the line in the middle is the median (Table S4).

**Supplemental Table S5. Description of experiments used in this work**

| **Figure** | **N° of plants** | **Treatment** | **Type of individual data** | **N° of individual data** | **Statistic test** |
| --- | --- | --- | --- | --- | --- |
| 1 A | 4 Col-0 | Mock (*Pst* DC3000) | Stomata | 193 | Tukey’s Method, p-value < 0.05 |
| 1 A | 4 Col-0 | *Pst* DC3000 1h (OD=0.1) | Stomata | 242 | Tukey’s Method, p-value < 0.05 |
| 1 A | 4 Col-0 | *Pst* DC3000 3h (OD=0.1) | Stomata | 233 | Tukey’s Method, p-value < 0.05 |
| 1 B | 3 Col-0 | Mock (*Pst* DC3000 *hrcC^-^*) | Stomata | 136 | Tukey’s Method, p-value < 0.05 |
| 1 B | 3 Col-0 | *Pst* DC3000 *hrcC^-^* 1h (OD=0.1) | Stomata | 160 | Tukey’s Method, p-value < 0.05 |
| 1 B | 3 Col-0 | *Pst* DC3000 *hrcC^-^* 3h (OD=0.1) | Stomata | 158 | Tukey’s Method, p-value < 0.05 |
| 1 A | 4 *des1* | Mock (*Pst* DC3000) | Stomata | 218 | Tukey’s Method, p-value < 0.05 |
| 1 A | 4 *des1* | *Pst* DC3000 1h (OD=0.1) | Stomata | 219 | Tukey’s Method, p-value < 0.05 |
| 1 A | 4 *des1* | *Pst* DC3000 3h (OD=0.1) | Stomata | 247 | Tukey’s Method, p-value < 0.05 |
| 1 B | 3 *des1* | Mock (*Pst* DC3000 *hrcC^-^*) | Stomata | 180 | Tukey’s Method, p-value < 0.05 |
| 1 B | 3 *des1* | *Pst* DC3000 *hrcC^-^* 1h (OD=0.1) | Stomata | 184 | Tukey’s Method, p-value < 0.05 |
| 1 B | 3 *des1* | *Pst* DC3000 *hrcC^-^* 3h (OD=0.1) | Stomata | 164 | Tukey’s Method, p-value < 0.05 |
| 2 A | 3 Col-0 | flg22 0 nM | Stomata | 134 | Tukey’s Method, p-value < 0.05 |
| 2 A | 3 Col-0 | flg22 10 nM | Stomata | 154 | Tukey’s Method, p-value < 0.05 |
| 2 A | 3 Col-0 | flg22 100 nM | Stomata | 112 | Tukey’s Method, p-value < 0.05 |
| 2 A | 3 Col-0 | flg22 1000 nM | Stomata | 158 | Tukey’s Method, p-value < 0.05 |
| 2 A | 3 *des1* | flg22 0 nM | Stomata | 130 | Tukey’s Method, p-value < 0.05 |
| 2 A | 3 *des1* | flg22 10 nM | Stomata | 125 | Tukey’s Method, p-value < 0.05 |
| 2 A | 3 *des1* | flg22 100 nM | Stomata | 140 | Tukey’s Method, p-value < 0.05 |
| 2 A | 3 *des1* | flg22 1000 nM | Stomata | 142 | Tukey’s Method, p-value < 0.05 |
| 2 B | 4 Col-0 | Control | Stomata | 278 | Tukey’s Method, p-value < 0.05 |
| 2 B | 4 Col-0 | flg22 1µM | Stomata | 283 | Tukey’s Method, p-value < 0.05 |
| 2 B | 4 Col-0 | GYY4137 100µM | Stomata | 229 | Tukey’s Method, p-value < 0.05 |
| 2 B | 4 Col-0 | flg22 + GYY4137 | Stomata | 207 | Tukey’s Method, p-value < 0.05 |
| 2 B | 4 *des1* | Control | Stomata | 256 | Tukey’s Method, p-value < 0.05 |
| 2 B | 4 *des1* | flg22 1µM | Stomata | 251 | Tukey’s Method, p-value < 0.05 |
| 2 B | 4 *des1* | GYY4137 100µM | Stomata | 236 | Tukey’s Method, p-value < 0.05 |
| 2 B | 4 *des1* | flg22 + GYY4137 | Stomata | 252 | Tukey’s Method, p-value < 0.05 |
| 3 | 4 Col-0 | *Pst* DC3000 *hrcC^-^* (OD=0.2) | Plant (4 leaf disc per pant) | 22 | Paired *t*-test, p-value 0.05 |
| 3 | 4 *des1* | *Pst* DC3000 *hrcC^-^* (OD=0.2) | Plant (4 leaf disc per pant) | 22 | Paired *t*-test, p-value 0.05 |
| 4 | 3 Col-0 | Control | Leaf discs | 36 | Not tested |
| 4 | 3 Col-0 | flg22 100 nM | Leaf discs | 36 | Unpaired t-test, p-value 0.01 |
| 4 | 3 *des1* | Control | Leaf discs | 36 | Not tested |
| 4 | 3 *des1* | flg22 100 nM | Leaf discs | 36 | Unpaired t-test, p-value 0.01 |
| 5 | 4 roGFP2-Orp1 | Mock | Guard cell | 77 | Tukey’s Method, p-value < 0.05;  Unpaired Wilcoxon, p-value < 0.05 |
| 5 | 4 roGFP2-Orp1 | flg22 1µM | Guard cell | 81 | Tukey’s Method, p-value < 0.05;  Unpaired Wilcoxon, p-value < 0.05 |
| 5 | 4 roGFP2-Orp1 | DTT 20 mM | Guard cell | 90 | Not tested |
| 5 | 4 roGFP2-Orp1 | H_2_O_2_ 10 mM | Guard cell | 78 | Not tested |
| 5 | 11 roGFP2-Orp1 + *des1* | Mock | Guard cell | 231 | Tukey’s Method, p-value < 0.05;  Unpaired Wilcoxon, p-value < 0.05 |
| 5 | 11 roGFP2-Orp1 + *des1* | flg22 1µM | Guard cell | 220 | Tukey’s Method, p-value < 0.05;  Unpaired Wilcoxon, p-value < 0.05 |
| 5 | 11 roGFP2-Orp1 + *des1* | DTT 20 mM | Guard cell | 211 | Not tested |
| 5 | 11 roGFP2-Orp1 + *des1* | H_2_O_2_ 10 mM | Guard cell | 225 | Not tested |
| 6 | 3 roGFP2-Orp1 | Mock | Guard cell | 113 | Tukey’s Method, p-value < 0.05 |
| 6 | 3 roGFP2-Orp1 | GYY4137 100µM | Guard cell | 103 | Tukey’s Method, p-value < 0.05 |
| 6 | 3 roGFP2-Orp1 | DTT 20 mM | Guard cell | 86 | Not tested |
| 6 | 3 roGFP2-Orp1 | H_2_O_2_ 10 mM | Guard cell | 66 | Not tested |
| 6 | 5 roGFP2-Orp1 x *rbohD* | Mock | Guard cell | 118 | Tukey’s Method, p-value < 0.05 |
| 6 | 5 roGFP2-Orp1 x *rbohD* | GYY4137 100µM | Guard cell | 227 | Tukey’s Method, p-value < 0.05 |
| 6 | 5 roGFP2-Orp1 x *rbohD* | DTT 20 mM | Guard cell | 113 | Not tested |
| 6 | 5 roGFP2-Orp1 x *rbohD* | H_2_O_2_ 10 mM | Guard cell | 92 | Not tested |
| 7 | 4 Col-0 | Control | Stomata | 422 | Tukey’s Method, p-value < 0.05 |
| 7 | 4 Col-0 | GYY4137 100 µM | Stomata | 363 | Tukey’s Method, p-value < 0.05 |
| 7 | 4 Col-0 | EGTA 200 µM | Stomata | 438 | Tukey’s Method, p-value < 0.05 |
| 7 | 4 Col-0 | BAPTA 25 µM | Stomata | 355 | Tukey’s Method, p-value < 0.05 |
| 7 | 4 Col-0 | GYY4137 + EGTA | Stomata | 396 | Tukey’s Method, p-value < 0.05 |
| 7 | 4 Col-0 | GYY4137 + BAPTA | Stomata | 300 | Tukey’s Method, p-value < 0.05 |
| S1 | 3 Col-0 | Control | Stomata | 240 | Tukey’s Method, p-value < 0.05 |
| S1 | 3 Col-0 | elf18 10 nM | Stomata | 197 | Tukey’s Method, p-value < 0.05 |
| S1 | 3 Col-0 | elf18 100 nM | Stomata | 212 | Tukey’s Method, p-value < 0.05 |
| S1 | 3 Col-0 | elf18 1000 nM | Stomata | 243 | Tukey’s Method, p-value < 0.05 |
| S1 | 3 *des1* | Control | Stomata | 224 | Tukey’s Method, p-value < 0.05 |
| S1 | 3 *des1* | elf18 10 nM | Stomata | 202 | Tukey’s Method, p-value < 0.05 |
| S1 | 3 *des1* | elf18 100 nM | Stomata | 247 | Tukey’s Method, p-value < 0.05 |
| S1 | 3 *des1* | elf18 1000 nM | Stomata | 246 | Tukey’s Method, p-value < 0.05 |
| 5 A | 4 Col-0 | Mock | Guard cell | 63 | Tukey’s Method, p-value < 0.05 |
| 5 A | 4 Col-0 | flg22 1µM | Guard cell | 76 | Tukey’s Method, p-value < 0.05 |
| 5 A | 4 Col-0 | flg22 1 µM + HT 200 µM | Guard cell | 70 | Tukey’s Method, p-value < 0.05 |
| 5 A | 4 Col-0 | DTT 20 mM | Guard cell | 69 | Not tested |
| 5 A | 4 Col-0 | H_2_O_2_ 10 mM | Guard cell | 76 | Not tested |
| S3 A and C | 3 replicates from 12 pooled Col-0 seedlings | flg22 0 min | seedling | 3 | Not tested |
| S3 A and C | 3 replicates from 12 pooled Col-0 seedlings | flg22 5 min | seedling | 3 | Not tested |
| S3 A and C | 3 replicates from 12 pooled Col-0 seedlings | flg22 15 min | seedling | 3 | Not tested |
| S3 A and C | 3 replicates from 12 pooled Col-0 seedlings | flg22 30 min | seedling | 3 | Not tested |
| S3 A and C | 3 replicates from 12 pooled *des1* seedlings | flg22 0 min | seedling | 3 | Not tested |
| S3 A and C | 3 replicates from 12 pooled *des1* seedlings | flg22 5 min | seedling | 3 | Not tested |
| S3 A and C | 3 replicates from 12 pooled *des1* seedlings | flg22 15 min | seedling | 3 | Not tested |
| S3 A and C | 3 replicates from 12 pooled *des1* seedlings | flg22 30 min | seedling | 3 | Not tested |
| S3 B | 3 replicates from 12 pooled Col-0 plants | Control | Epidermal peels (GCe-RNA) | 3 | t-test, p-value < 0.0001 |
| S3 B | 3 replicates from 12 pooled Col-0 plants | flg22 1 µM | Epidermal peels (GCe-RNA) | 3 | t-test, p-value < 0.0001 |
| S3 B | 3 replicates from 12 pooled *des1* plants | Control | Epidermal peels (GCe-RNA) | 3 | t-test, p-value < 0.0001 |
| S3 B | 3 replicates from 12 pooled *des1* plants | flg22 1 µM | Epidermal peels (GCe-RNA) | 3 | t-test, p-value < 0.0001 |
| S3 D | 3 replicates from 12 pooled Col-0 plants | Control | Epidermal peels (GCe-RNA) | 3 | t-test, p-value < 0.0001 |
| S3 D | 3 replicates from 12 pooled Col-0 plants | flg22 1 µM | Epidermal peels (GCe-RNA) | 3 | t-test, p-value < 0.0001 |
| S3 D | 3 replicates from 12 pooled *des1* plants | Control | Epidermal peels (GCe-RNA) | 3 | t-test, p-value < 0.0001 |
| S3 D | 3 replicates from 12 pooled *des1* plants | flg22 1 µM | Epidermal peels (GCe-RNA) | 3 | t-test, p-value < 0.0001 |
| S4 | 4 replicates from 12 pooled Col-0 plants | Control | Epidermal peels (GCe-RNA) | 4 | t-test, p-value < 0.0001 |
| S4 | 4 replicates from 12 pooled Col-0 plants | flg22 1 µM | Epidermal peels (GCe-RNA) | 4 | t-test, p-value < 0.0001 |

**Supplemental Table S6. Primer sequence used in this work**

| **Name** | **Sequence** | **Gene** | **Reference** | **Experiment** |
| --- | --- | --- | --- | --- |
| qACT2_Fw | GCCATCCAAGCTGTTCTCTC | At3g18780 | (Soto et al., 2015) | RT-qPCR |
| qACT2_Rv | GAAACCCTCGTAGATTGGCA | At3g18780 | (Soto et al., 2015) | RT-qPCR |
| qDES1_Fw | GAAGCTGCGAATTTGCCAGTTG | At5g28030 | This work | RT-qPCR |
| qDES1_Rv | AAATGTAACCTTGGTACCAAC | At5g28030 | This work | RT-qPCR |
| qWRKY22_Fw | GATCATCTAGCGGTGGGAGA | At4g01250 | (Macho et al., 2012) | RT-qPCR |
| qWRKY22_Rv | TATTCCTCCGGTGGTAGTGG | At4g01250 | (Macho et al., 2012) | RT-qPCR |
| qFRK1_Fw | ATCTTCGCTTGGAGCTTCTC | At2g19190 | (He et al., 2006) | RT-qPCR |
| qFRK1_Rv | TGCAGCGCAAGGACTAGAG | At2g19190 | (He et al., 2006) | RT-qPCR |
| DES1_Fw | CAAAGGATTGATTACTCCGGGAAAG | At5g28030 | This work | Genotyping |
| DES1_Rv | CTGGAATTTCACCAGAGCCTATTC | At5g28030 | This work | Genotyping |
| LB | TGGTTCACGTAGTGGGCCATCG | At5g28030 | This work | Genotyping |
| rbohD_211 | GTCGCCAAAGGAGGCGCCGA | At5g47910 | (Torres et al., 2002) | Genotyping |
| rbohD_92b | GGATACTGATCATAGGCGTGGCTCCA | At5g47910 | (Torres et al., 2002) | Genotyping |
| dSpm1 | CTTATTTCAGTAAGAGTGTGGGGTTTTGG | At5g47910 | (Torres et al., 2002) | Genotyping |
